# Therapeutic targeting inflammation linking periodontitis and atherosclerotic comorbidities using cell-free DNA-capturing nanomaterials

**DOI:** 10.64898/2026.03.01.708715

**Authors:** Hanyao Huang, Xuanzhi Zhu, Xiao Chen, Fangman Chen, Shouzheng Cheng, Suwan Ding, Yang Xiao, Xiaochun Xie, Chuanxu Cheng, Renjie Yang, Jiali Chen, Jing Liu, Xiaoming Yang, Chao Yang, Bing Shi, Dan Shao, Lei Zhao, Kam W. Leong

**Affiliations:** State Key Laboratory of Oral Diseases and National Clinical Research Center for Oral Diseases and Department of Oral and Maxillofacial Surgery, West China Hospital of Stomatology, Sichuan University, Chengdu, Sichuan 610041, China; Department of Biomedical Engineering, Columbia University, New York, NY 10027, USA; State Key Laboratory of Oral Diseases and National Clinical Research Center for Oral Diseases and Department of Periodontics, West China Hospital of Stomatology, Sichuan University, Chengdu, Sichuan 610041, China; National Engineering Research Center for Tissue Restoration and Reconstruction, South China University of Technology, Guangzhou, Guangdong, 510006, China; State Key Laboratory of Oral Diseases and National Clinical Research Center for Oral Diseases and Eastern Clinic, West China Hospital of Stomatology, Sichuan University, Chengdu, Sichuan 610041, China; West China School of Public Health and West China Fourth Hospital, Sichuan University, Chengdu, 610041, Sichuan, China; Department of Orthopedics, Guangdong Provincial Key Laboratory of Bone and Joint Degeneration Diseases, The Third Affiliated Hospital of Southern Medical University, Guangzhou, Guangdong 510630, China; School of Medicine, South China University of Technology, Guangzhou, Guangdong 510006, China; School of Biomedical Sciences and Engineering, Guangzhou International Campus, South China University of Technology, Guangzhou, Guangdong 510006, China; Department of Systems Biology, Columbia University Medical Center, New York, NY 10032, USA

## Abstract

Periodontitis-associated systemic inflammation makes it a great challenge to explore therapeutic options applicable to periodontitis and atherosclerotic comorbidities. Here, we identify the crucial role of cell-free DNA (cfDNA) that underlies these comorbidities. Hypothesizing cfDNA as a therapeutic target, we engineer polyamidoamine dendrimer-functionalized nanomaterials to modulate such local-systemic inflammatory crosstalk. Periodontium-originated DNA can be systemically captured by cationic nanomaterials, and capturing cfDNA, whether locally or systemically, alleviates both periodontitis and atherosclerosis prior to severe atherosclerotic development *in vivo*. The transcriptomic and single-cell RNA sequencing analyses together reveal that cfDNA-capturing nanomaterials regulate inflammatory foam cell transformation in macrophages by modulating the expression of lipid-related foamy markers *Spp1* and *Fabp4*. This study provides a proof of concept for cfDNA-driven periodontitis-atherosclerosis crosstalk, and offers a cfDNA-capturing nanoplatform for therapeutic intervention targeting periodontitis and atherosclerotic comorbidities in a holistic fashion.

## Introduction

The interaction between local and systemic inflammation and the treatment of such inflammatory comorbidities remains a significant challenge [1]. Periodontitis, a common local inflammatory disease that poses a significant public health burden, involves microbe-induced inflammatory bone loss and destruction of periodontal tissue [2], and it is linked to various systemic diseases [3]. On the other hand, atherosclerosis, one of the most prevalent systemic inflammatory conditions, is the leading cause of mortality worldwide [4]. It is characterized by chronic vascular inflammation, leading to lipid accumulation and the development of fibro-lipid structures known as atherosclerotic plaques [5]. Evidence indicates that periodontitis is an independent risk factor for atherosclerosis [6–9]. However, it has been largely uncertain by which this imbalanced relationship can causally link periodontitis and atherosclerosis. From a medical and therapeutic standpoint, it is imperative to understand whether the periodontitis-atherosclerosis crosstalk is merely a correlative one or orchestrated by causal mechanistic interactions.

Since the progression of periodontitis and atherosclerosis is driven by a proinflammatory immune response, inappropriate activation of the immune system may be a key factor in the development and crosstalk between these local-systemic inflammatory comorbidities [3, 10, 11]. Pattern-recognition receptors, such as toll-like receptors (TLRs), which respond to molecular patterns, can be implicated in the crosstalk between periodontitis and atherosclerosis [12]. TLR9, in particular, recognizes cell-free DNA (cfDNA) from damaged host cells, as well as from external bacteria or viruses [13]. The role of the cfDNA-TLR9-initiated and -mediated proinflammatory immune response has already been explicated in the pathogenesis of both periodontitis and atherosclerosis [14–17]. Pathogenic DNA from periodontitis has been detected in the circulation and atherosclerotic plaques [18, 19], indicating that cfDNA may originate from periodontitis and contribute to the periodontitis-associated progression of atherosclerosis [20]. Thus, it is reasonable to hypothesize that cfDNA acts as a messenger to drive the crosstalk between these two diseases. Targeting the cfDNA-mediated inflammation may offer a therapeutic approach for managing these local-systemic inflammatory comorbidities.

In recent years, DNA-targeting immunomodulating therapies based on biomaterials have made significant strides, showing therapeutic efficacy in various inflammatory conditions [21–31]. One promising approach involves nanomaterials-based cfDNA capturing strategies, which have demonstrated notable potential in treating circulation-related diseases by inhibiting cfDNA-mediated inflammation, as seen in cases like sepsis [26, 29]. Here, we propose the design of cfDNA-targeting nanomaterials to act as an immunomodulator, aimed to prove the concept that capturing cfDNA can control the crosstalk between periodontitis and atherosclerosis. In this study, we decipher the role of cfDNA in the crosstalk between periodontitis and atherosclerosis, investigate the potential of functional nanomaterials in capturing cfDNA that driven by local inflammation in circulation, and demonstrate the therapeutic efficacy and mechanisms of these nanomaterials in controlling local-systemic inflammatory comorbidities.

## Results

### Plasma cfDNA levels correlate with periodontitis severity but show no correlation with atherosclerosis severity

We employ a cross-sectional study design to measure the concentrations of cfDNA in plasma among three groups of individuals following strict inclusion criteria (77 subjects in total), including the healthy group (25 healthy volunteers), the periodontitis group (27 patients with periodontitis but without atherosclerosis), and the periodontitis plus atherosclerosis group (25 patients with both periodontitis and atherosclerosis) (Fig. 1c). No atherosclerosis patients with healthy periodontal condition are observed during the sample collection. Detailed demographic and clinical data for all study participants are provided in Supplementary Table 1. The correlations between cfDNA and clinical parameters of periodontitis and atherosclerosis are analyzed.

**Fig. 1.**
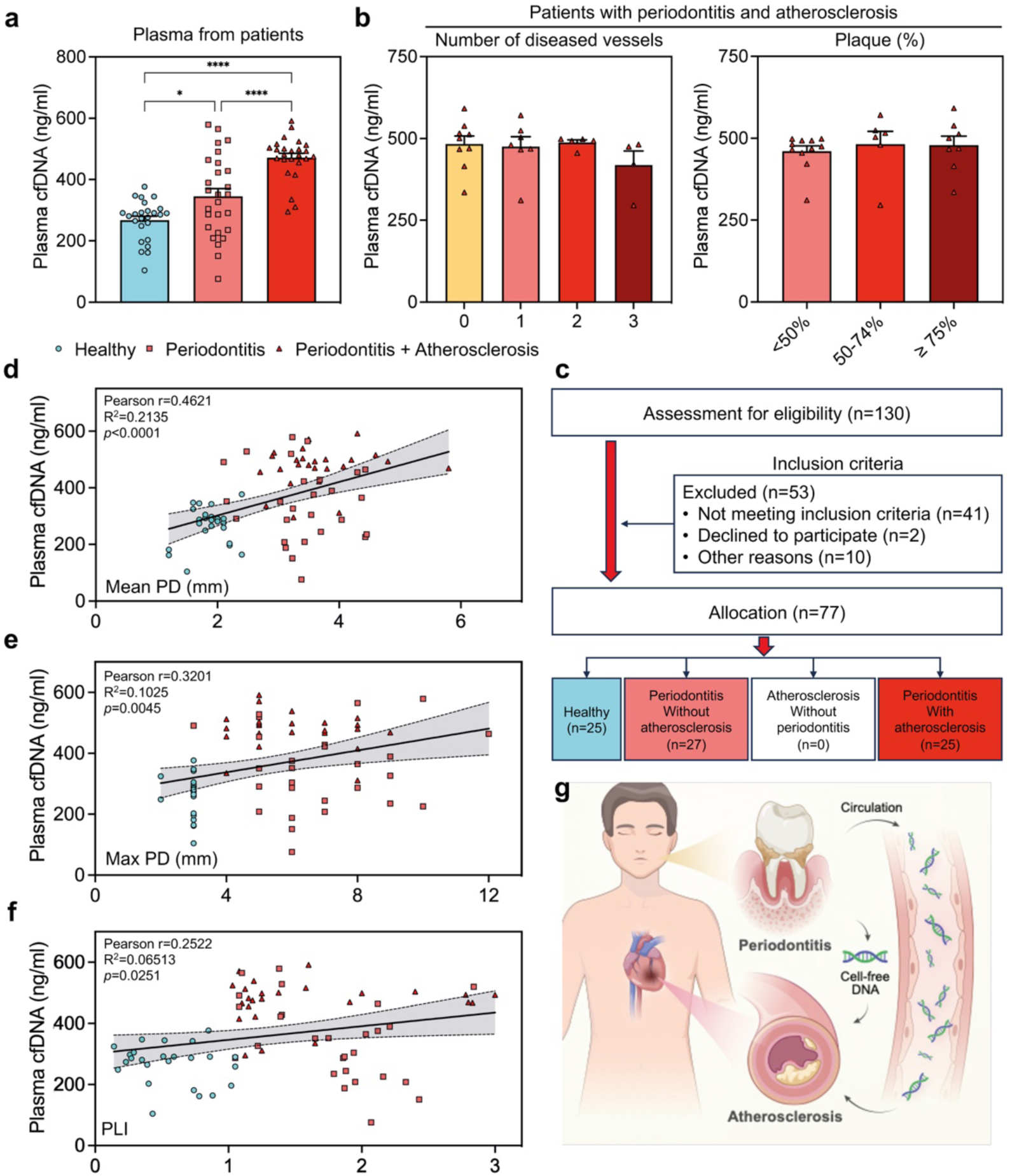
Plasma cfDNA concentration is significantly correlated with periodontal clinical parameters, but has a weak correlation with atherosclerosis clinical parameters. **a** cfDNA levels in plasma of healthy volunteers (n = 25), periodontitis without atherosclerosis patients (n = 27) and periodontitis patients with atherosclerosis (n = 25). **b** Plasma cfDNA levels in subgroups with different numbers of diseased vessels and different percentage of plaque area in the lumen in patients with periodontitis and atherosclerosis. Data are means ± SEM; differences were assessed by one-way analysis of variance and Tukey’s multiple comparisons test. *P < 0.05, **P < 0.01, ***P < 0.001, ****P < 0.0001. **c** Cross-sectional study design inclusion and exclusion flowchart. **d-f** Scatter plot of plasma cfDNA levels and parameters of periodontal examination. Pearson correlation analysis and linear regression analysis were conducted (n = 77). The black line is the fitted regression line, and the light grey shading around it is the 95% confidence interval. PD: probing depth (mm); PLI: plaque index (1-3). **g** Schematic diagram of the association between periodontitis and atherosclerosis through circulating cell-free DNA.

Plasma cfDNA levels are significantly higher in patients with periodontitis and in those with both periodontitis and atherosclerosis compared to healthy volunteers, with the highest levels observed in patients with both periodontitis and atherosclerosis (Fig. 1a). In patients with periodontitis and atherosclerosis, there are no significant differences in plasma cfDNA concentrations between different numbers of diseased vessels, the percentage of plaque area in the lumen, and probing depth (PD) (Fig. 1b and Supplementary Fig. 1). However, there are significant positive correlations between plasma cfDNA and periodontal clinical parameters, including mean probing depth (PD), maximum PD, and plaque index (PI) (Fig. 1d-f, Supplementary Table 2). These results demonstrate that cfDNA can play a critical role in periodontitis-atherosclerosis crosstalk (Fig. 1g).

### Periodontium-originated DNA has multi-organ distribution and be captured by systemic-administrated functional nanomaterials

The strategy to capture negatively charged biomacromolecules by cationic nanomaterials has shown great potential in treating inflammatory diseases [25]. Here, we use third-generation polyamidoamine dendrimer (PG3)-functionalized mesoporous silica nanoparticles (MSNs) (PG3@MSNs) to explicate the messaging role of cfDNA in crosstalk between periodontitis and atherosclerosis. The pristine MSNs with an average diameter of approximately 50 nm have a high surface area (664.38 m² g^-1^) and large pore volume (1.38 cm³ g^-1^) with a narrow pore size distribution centered at 8.31 nm (Fig. 2a-b and Supplementary Fig. 2a-c). The PG3@MSNs are fabricated by conjugating PG3 to MSNs, imparting a positively charged surface without altering the morphology or size of MSNs (Fig. 2b-d and Supplementary Fig. S2d). Free PG3 and PG3@MSNs exhibit a high binding affinity for calf thymus DNA (ct-DNA), whereas unmodified MSNs do not (Fig. 2e). Though its DNA binding affinity is slightly lower than free PG3, PG3@MSNs cause significantly less cytotoxicity in cells (Fig. 2f and Supplementary Fig. 2e). We then evaluate whether PG3@MSNs can capture cfDNA and inhibit inflammation. Both PG3@MSNs and free PG3 significantly inhibit CpG DNA-induced TLR9 activation, reducing TNF-α and IL-6 secretion by macrophages (Fig. 2g-h and Supplementary Fig. 2f). Additionally, PG3@MSNs exhibit H_2_O_2_-responsive degradation (Supplementary Fig. S3).

**Fig. 2.**
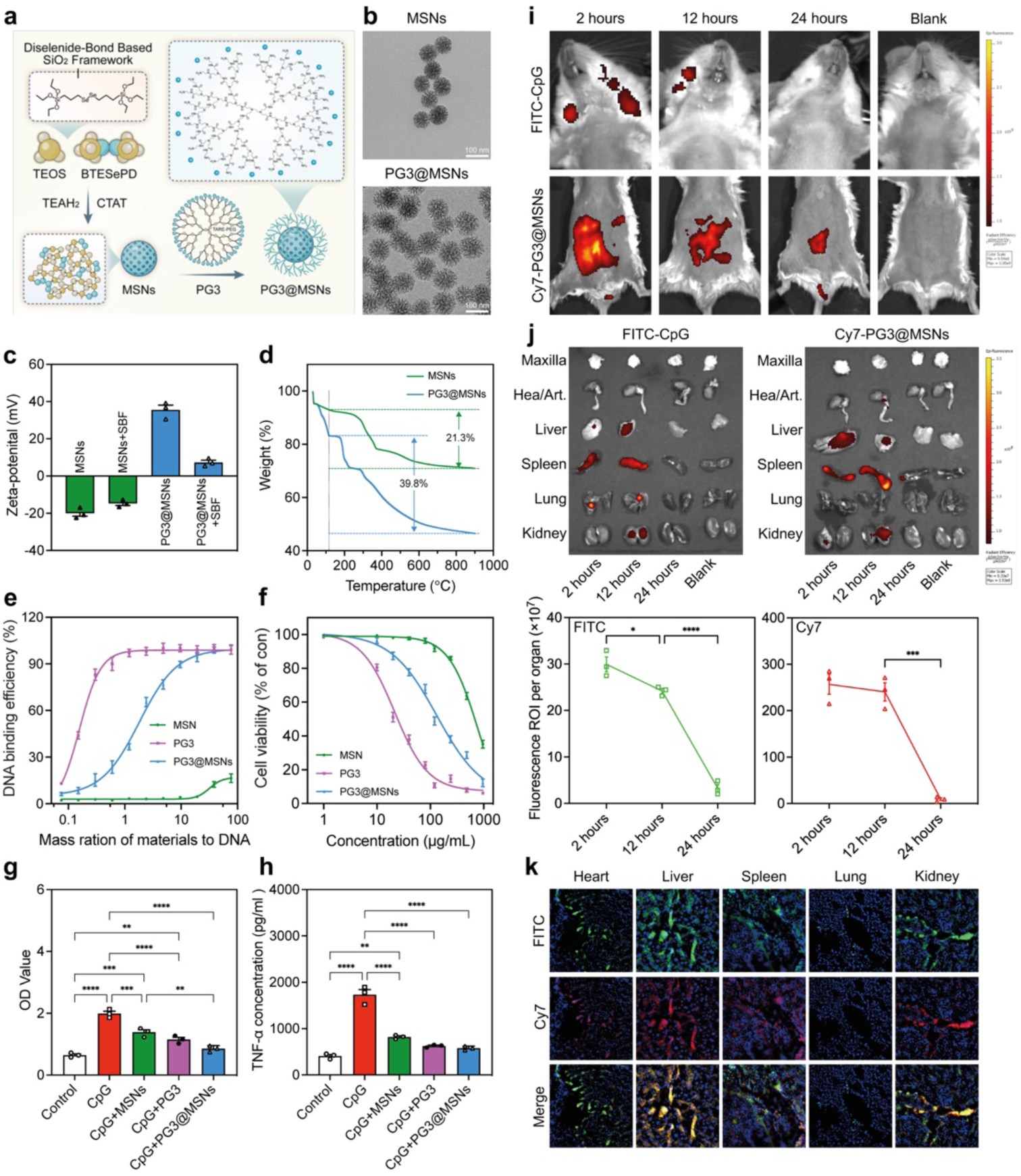
The synthesis, characterization, in vitro activity, and in vivo distribution of PG3@MSNs. **a** The schematic representation of the design and synthesis of PG3@MSNs. **b** TEM imaging of MSNs before and after coating with PG3. Scale bar, 100 nm. **c** Zeta potential of MSNs and PG3@MSNs with of without SBF (n = 3). SBF, simulated body fluids. **d** The thermal decomposition rate of the nanoparticles changes after coating with PG3. **e** DNA binding efficiency of MSNs, PG3, and PG3@MSNs at different nanoparticle:DNA mass ratios at 37°C. **f** Viability of RAW264.7 macrophages treated for 24 hours with various concentrations of MSNs, PG3, and PG3@MSNs. **g** Activation of HEK-Blue TLR9 reporter cells by CpG DNA (ODN 1826) in the absence or presence of MSNs, PG3, and PG3@MSNs for 24 hours. The corresponding SEAP activity in supernatants from each group was determined with a QUANTI-Blue assay at OD_620_. **h** RAW 264.7 macrophages were stimulated with CpG DNA (ODN 1826) in the absence or presence of MSNs, PG3, and PG3@MSNs for 24 hours. Supernatants were assayed for TNF-α by ELISA. **i, j** In vivo (**i**) and ex vivo (**j**) imaging of FITC-CpG DNA (periodontal injection) and Cy7-PG3@MSNs (i.p. injection) signal at 2, 12, 24 hours. PBS-treated (blank) mice were used as the basal reference. Semi-quantitative analysis of ex vivo fluorescence images of the organs was conducted (n =3). Data are means ± SEM; differences were assessed by one-way analysis of variance and Tukey’s multiple comparisons test. \**P* < 0.05, \*\**P* < 0.01, \*\*\**P* < 0.001, \*\*\*\**P* < 0.0001. ROI, region of interest. **k** Immunofluorescence staining reveals the co-localization of FITC-CpG DNA (periodontal injection) and Cy7-PG3@MSNs (i.p. injection) within the vasculature of major organs. Scale bar, 100 μm.

To verify whether DNA originating from the periodontium can be cleared by systemically administered biomaterials, we locally inject FITC-labeled CpG DNA into the periodontal tissues of mice while concurrently administering Cy7-labeled PG3@MSNs via intraperitoneal injection. Using the in vivo imaging system (IVIS), FITC signals are detectable locally in the mouse periodontium at 2 and 12 hours post-injection, while Cy7 signals are detectable at 2, 12, and 24 hours post-injection (Fig. 2i). Within 24 hours, both the periodontally injected FITC-CpG and intraperitoneally injected Cy7-PG3@MSNs are distributed in major organs such as the heart, liver, spleen, lung, and kidney, with signal intensities decreasing over time (Fig. 2j). Immunofluorescence staining reveals co-localization of FITC and Cy7 signals in these organs, indicating that DNA originating from the periodontium can successfully be captured by intraperitoneally injected biomaterials in the circulation (Fig. 2k).

### Systemically capturing cfDNA alleviates both atherosclerosis and periodontitis

Firstly, we aim to determine the impact of systemically capturing cfDNA on a comorbidity model, ApoE^-/-^ mice with ligature-induced periodontitis and western diet-induced atherosclerosis (Fig. 3a and b). Four groups are designed: control group without ligature (Control), untreated group with ligature and phosphate-buffered saline (PBS) injection (Ligature), PG3@MSNs treated group with ligature and PG3@MSNs injection (Ligature + PG3@MSNs), and a PG3@MSNs treated group without ligature (PG3@MSNs). All mice receive intraperitoneal injections of PBS or PG3@MSNs on alternate days after ligature completion, and no notable toxicity is observed (Fig. 3c and Supplementary Fig. 4).

**Fig. 3.**
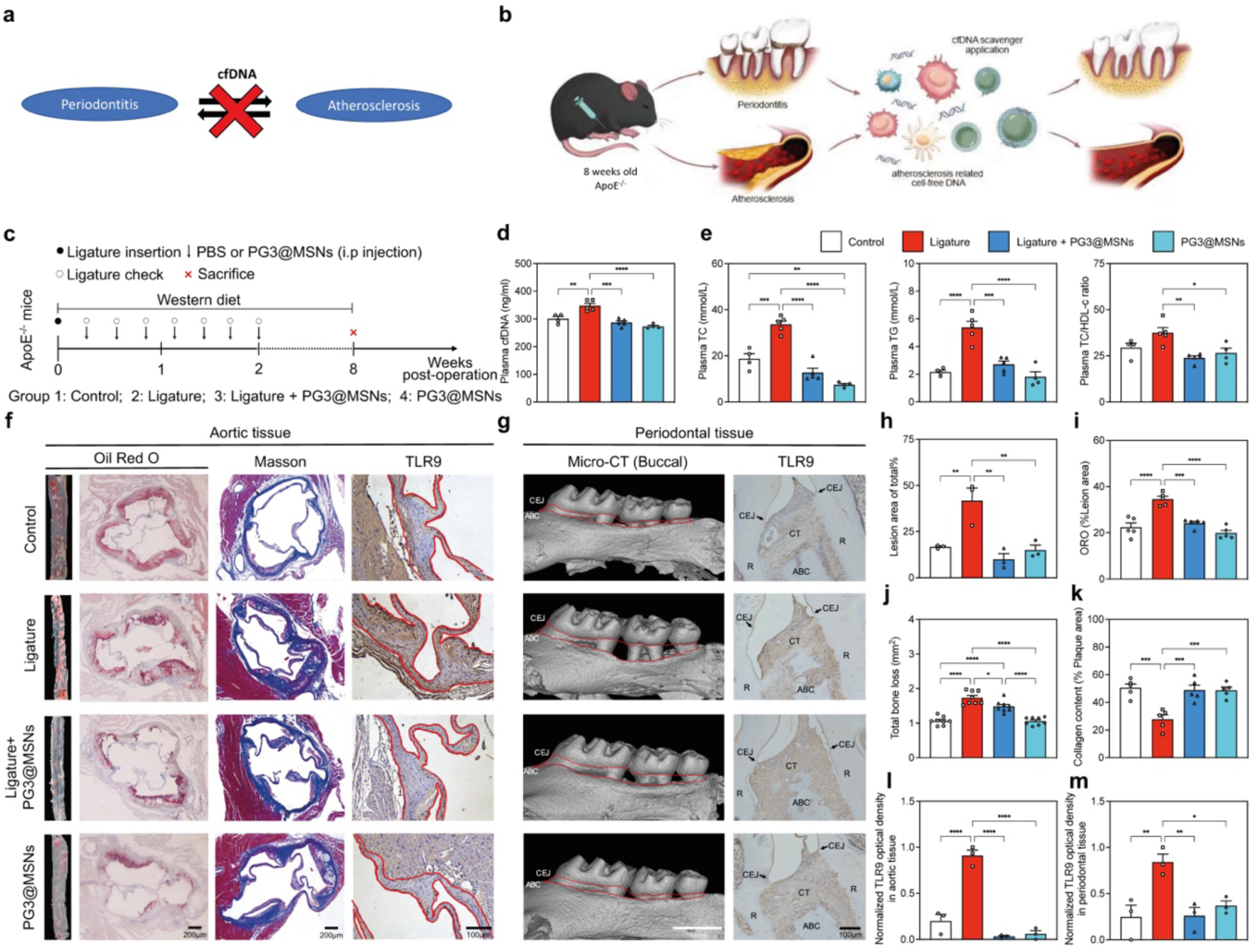
Intraperitoneal injection of PG3@MSNs significantly attenuates ligature-induced periodontal alveolar bone resorption and concurrently decelerates the progression of atherosclerotic plaque formation in ApoE^-/-^ mice. **a**, **b** Schematic illustration of the comorbidity relationship and the mechanism of PG3@MSNs systemically application. **c** Experimental schedule of in vivo study. **d, e** Plasma cfDNA (**d**), TC, TG concentration, and the TC to HDL-c ratio (**e**) at 8 weeks post operation (n = 4-5). TC, total cholesterol. TG, total triglycerides. HDL-c, high-density lipoprotein cholesterol. **f** Gross and histopathological staining of the thoracoabdominal aorta and aortic valve tissues. The columns from left to right represent gross Oil Red O staining of the thoracoabdominal aorta, Oil Red O staining, Masson’s trichrome staining (scale bar, 200 μm), and TLR9 immunohistochemical staining (Scale bar, 100 μm) of the aortic valve. **g** Periodontal tissue Micro-CT buccal view (left column; scale bar, 1mm) and TLR9 immunohistochemical staining (right column; scale bar, 100 μm). CEJ, cemento-enamel junction. ABC, alveolar bone crest. CT, connective tissue. R, root. **h, i** Quantification of Oil Red O staining in thoracoabdominal aorta (**h**) (n = 3) and aortic valve (**i**) (n = 5). **j** Quantification of total alveolar bone resorption (n = 8). **k** Quantification of collagen content within atherosclerotic plaques (n = 5). **l, m** Expression of TLR9 in the aortic valve (**l**) and epithelium of periodontal tissues (**m**) (n = 3). Data are means ± SEM; differences were assessed by one-way analysis of variance and Tukey’s multiple comparisons test. \**P* < 0.05, \*\**P* < 0.01, \*\*\**P* < 0.001, \*\*\*\**P* < 0.0001.

The ligature group has elevated plasma cfDNA, total cholesterol (TC), total triglycerides (TG) concentration, and the TC to high-density lipoprotein cholesterol (HDL-c) ratio, as well as plasma proinflammatory cytokines, including tumor necrosis factor-α (TNF-α) and interleukin-6 (IL-6) (Fig. 3d and e, and Supplementary Fig. 5). Severe lipid deposition and enlarged area of the plaques in the lumen, as well as alveolar bone loss in periodontal tissue are found in the ligature group (Fig. 3f-j, and Supplementary Fig. 6-8). Masson staining reveals a decreased proportion of fiber content within the plaques, indicating increased plaque instability (Fig. 3f and k). TLR9 expression and TLR9 pathway-related mRNA epression and cytokines in both aortic and periodontal tissues are upregulated in the ligature group (Fig. 3f, g, l, and m, and Supplementary Fig. 9 and 10). Cellular lipid uptake-related mRNA expression, including *Sr-a1* and *Acat1*, in aortic tissue are also upregulated in the ligature group, along with downregulation of the cellular lipid efflux gene *Abca1* (Supplementary Fig. 11).

Systemically-administrated PG3@MSNs significantly decrease the levels of plasma cfDNA, TC, TG, and TC/HDL-c ratio, as well as TNF-α and IL-6 (Fig. 3d and e, and Supplementary Fig. 5). Most interestingly, systemic administration of PG3@MSNs can alleviate both alveolar bone loss and arterial plaques by reducing TLR9-mediated proinflammatory response while controlling lipid metabolism (Fig. 3d-m, and Supplementary Fig. 5-11). *In vitro* study demonstrates that plasma samples from the ligature group induce a significantly stronger inflammation to HEK-TLR9 reporter cells and RAW 264.7 cells, which can be alleviated by PG3@MSNs (Supplementary Fig. 12). These results suggest that experimental periodontitis can significantly promote the progression of atherosclerosis, and the nanomaterial-functionalized cfDNA capturing strategy can inhibit this promotion and potentially block the crosstalk between these local-systemic inflammatory comorbidities.

### Locally blocking cfDNA from periodontal tissue alleviated atherosclerosis

To further confirm that cfDNA capturing can help block the crosstalk between these local-systemic inflammatory comorbidities, we choose to control the local originating of cfDNA and test whether the periodontal administration of PG3@MSNs affects the progression of atherosclerosis. The animal model establishment follows the same approach as the previously described PG3@MSNs systemically-administered experiments, with the only modification being the local administration of PG3@MSNs around the ligatured molars instead of through systemic administration (Fig. 4 a-c) [27]. Similar to PG3@MSNs systemically-administered experiments, the ligature group has elevated levels of plasma cfDNA, TC, TG, and TC/HDL-c ratio, and locally-administered PG3@MSNs can reduce those levels (Fig. 4d and e), and can alleviate ligature-induced alveolar bone loss in periodontal tissue, as well as severe lipid deposition and enlarged area of the plaques in the lumen (Fig. 4f-j and Supplementary Fig. 13-15). Ligature-induced arterial plaque instability is also preserved by locally-administered PG3@MSNs (Fig. 4f and k). Ligature-induced upregulation of TLR9 expression and TLR9 pathway-related mRNA and cytokines in both aortic and periodontal tissues can be alleviated by locally-administered PG3@MSNs (Fig. 4f, g, l and m, and Supplementary Fig. 16-18). Meanwhile, locally-administered PG3@MSNs can reverse the ligature-induced changes in lipid metabolism-related mRNA expression (Supplementary Fig. 19).

**Fig. 4.**
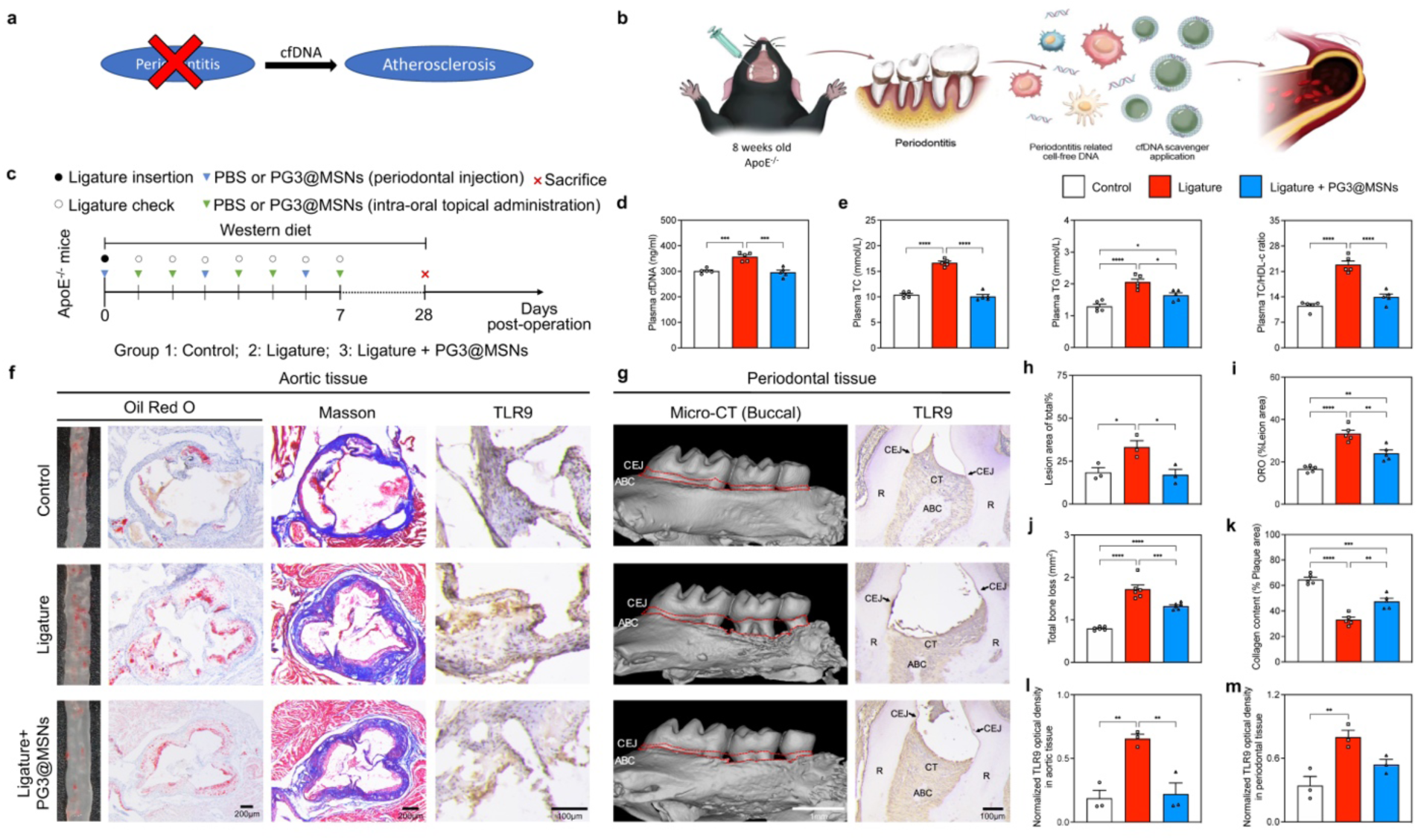
Local injection of PG3@MSNs attenuates both ligature-induced periodontal alveolar bone resorption and atherosclerotic plaque formation in ApoE^-/-^ mice. **a**, **b** Schematic illustration of the comorbidity relationship and the mechanism of PG3@MSNs locally application. **c** Experimental schedule of in vivo study. **d, e** Plasma cfDNA (**d**), TC, TG concentration, and the TC to HDL-c ratio (**e**) at 4weeks post operation (n = 5). TC, total cholesterol. TG, total triglycerides. HDL-c, high-density lipoprotein cholesterol. **f** Representative images of histopathological staining of the thoracoabdominal aorta and aortic valve tissues. The columns from left to right represent gross Oil Red O staining of the thoracoabdominal aorta, Oil Red O staining, Masson’s trichrome staining (scale bars, 200 μm), and TLR9 immunohistochemical staining (scale bar, 100 μm) of the aortic valve. **g** Representative images of 3D reconstruction of mice maxilla (left; scale bar, 1mm) and TLR9 immunohistochemical staining (right; scale bar, 100 μm). CEJ, cemento-enamel junction. ABC, alveolar bone crest. CT, connective tissue. R, root. **h, i** Quantification of Oil Red O staining in thoracoabdominal aorta (**h**) (n = 3) and aortic valve (**i**) (n = 5). **j** Quantification of total alveolar bone resorption (n = 8). **k** Quantification of collagen content within atherosclerotic plaques (n = 5). **l, m** Expression of TLR9 in the aortic valve (**l**) and epithelium of periodontal tissues (**m**) (n = 3). Data are means ± SEM; statistical analysis was performed using one-way ANOVA with Tukey’s post hoc test; \**P* < 0.05, \*\**P* < 0.01, \*\*\**P* < 0.001, \*\*\*\**P* < 0.0001.

Then, we test whether controlling the systemic originating of cfDNA affects the progression of periodontitis. Severe atherosclerosis in ApoE^-/-^ mice is induced by a 3-month western diet, followed by a two-week systemic administration of PG3@MSNs, and then ligature placement to induce periodontitis (Supplementary Fig. 20a and b). By comparison with the wide-type mice with periodontitis, atherosclerotic mice encountered more damage on supportive periodontium tissue loss, which cannot be alleviated by systemically-administrated PG3@MSNs (Supplementary Fig. 20 c, d), suggesting that atherosclerosis deteriorated the experimental periodontitis. Herein, we confirm that systemically- and locally-administrated cfDNA capturing strategy can inhibit the progression of local-systemic inflammatory comorbidities before systemic inflammation establishment.

### Capturing cfDNA manipulates lipid-related foamy change of macrophages

Cholesterol-laden macrophages, also known as foamy cells, play a pivotal role in atherosclerosis pathogenesis [32]. Our previous data suggest that plasma cfDNA levels are indicative of inflammation severity and may be associated with lipid metabolism (Fig. 3d, e and Fig. 4d, e). Therefore, we aim to investigate how cfDNA-capturing nanomaterials simultaneously affect the proinflammatory response and lipid metabolism in macrophages at the same time. RAW 264.7 macrophages are exposed to oxidized low-density lipoprotein (oxLDL) and CpG DNA *in vitro* to simulate the conditions found in mice with both periodontitis and atherosclerosis, with varying treatments including DNase I, PG3, and PG3@MSNs (Fig. 5a). Our findings indicate that CpG DNA amplifies oxLDL-induced cytoplasmic lipid accumulation and markedly elevates levels of total cholesterol (TC), free cholestorl (FC), cholesterol esters (CE), and the CE/TC rate, as well as proinflammatory cytokine secretion in RAW 264.7 cells (Fig. 5b-e). Both PG3 and PG3@MSN effectively reverse these changes, while DNase I application produces less consistent results (Fig. 5b-e).

**Fig. 5.**
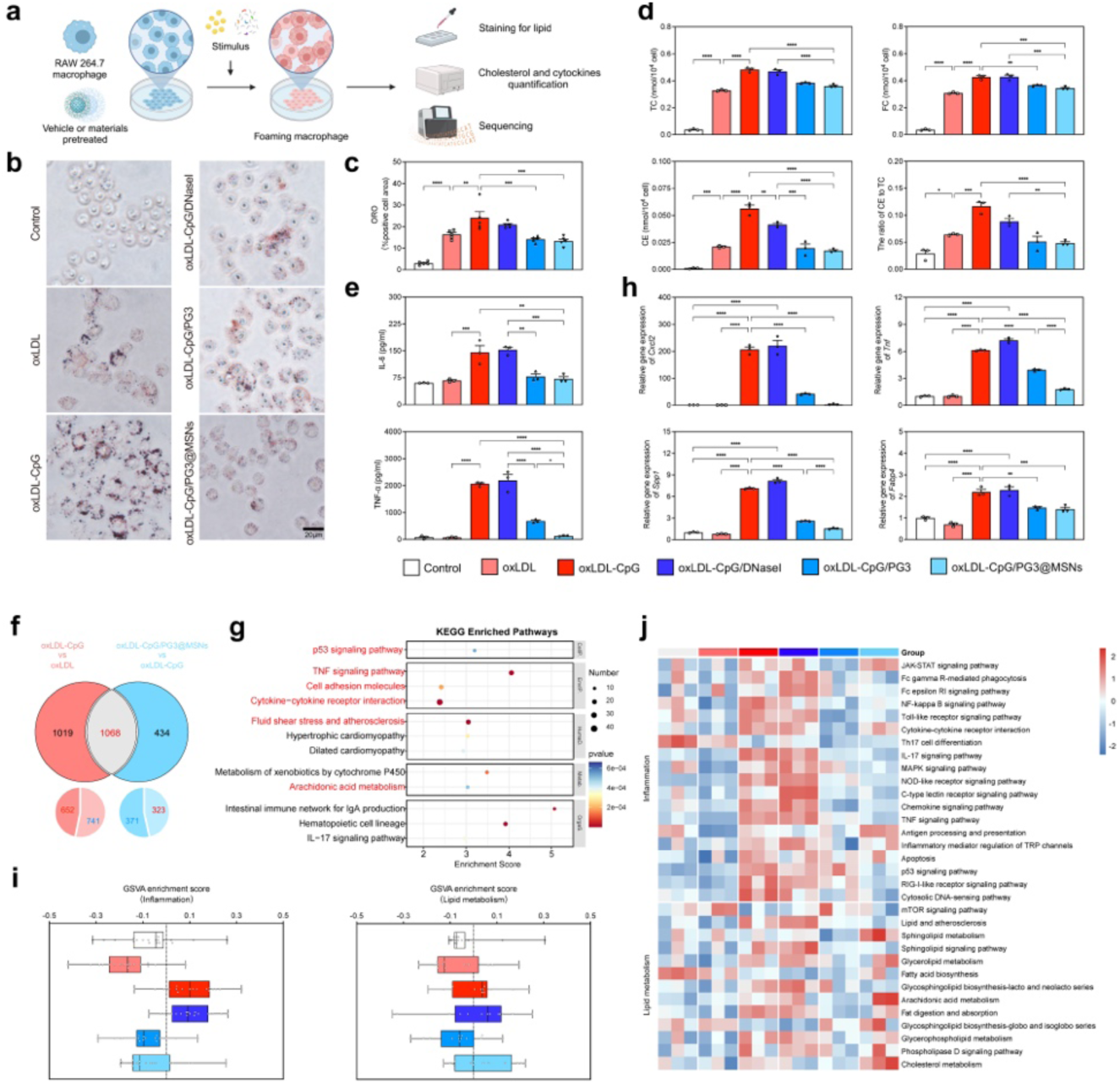
PG3@MSNs attenuates cfDNA-manipulated foamy change in macrophages. **a** Schematic of the design of the in vitro models. **b** Representative images of Oil Red O staining of RAW 264.7 macrophages. Scar bar, 20 μm. **c** Quantification of Oil Red O staining of RAW 264.7 macrophages in **b** (n = 5). **d** TC, FC, and CE concentration, and the CE to TC ratio of RAW 264.7 macrophages in **a** (n = 3). TC, total cholesterol; FC, free cholesterol; CE, cholesteryl ester. **e** Quantification of supernatant IL-6 and TNF-α concentrations using ELISA (n = 3). **f** Venn diagram of the intersection of differentially expressed genes. **g** Enriched KEGG pathways of the intersection genes in **f**. **h** Relative gene expression of *Cxcl2*, *Tnf*, *Spp1*, and *Fabp4* in RAW 264.7 macrophages in **a**. **i** Gene set variation analysis (GSVA) score of inflammation, and lipid metabolism-related pathways of RAW 264.7 macrophages. **j** Heatmap showing the GSVA-enriched pathways related to inflammation and lipid metabolism in **i**. Data in **c-d**, and **h** are means ± SEM; statistical analysis was performed using one-way ANOVA with Tukey’s post hoc test (**c-d**, **h**); \**P* < 0.05, \*\**P* < 0.01, \*\*\**P* < 0.001, \*\*\*\**P* < 0.0001.

Additionally, transcriptomic analysis is conducted to further validate the therapeutic benefits of cationic materials in reducing inflammation and regulating lipid metabolism in RAW 264.7 cells. Differentially expressed genes (DEGs) are identified between each pair of groups (Supplementary Fig. 21a, b). Compared to DNase I and PG3, PG3@MSNs demonstrate the greatest protective effects on macrophages against oxLDL-CpG DNA challenge, reversing 741 of 1393 upregulated genes and 323 of 694 downregulated genes induced by CpG DNA, leading to 1068 genes for further investigation (Fig. 5f and Supplementary Fig. 21c). Kyoto Encyclopedia of Genes and Genomes (KEGG) pathway analysis demonstrates that the differentially expressed genes after oxLDL-CpG DNA challenge and PG3@MSNs treatment are involved in inflammation, lipid metabolism, and cardiovascular disease (Fig. 5g and Supplementary Fig. 21d). The addition of CpG DNA induces a broad upregulation of inflammatory and foam cell-associated genes in macrophages (Fig. 5h and Supplementary Fig. 22) [33, 34]. Gene set enrichment analysis (GSEA) shows upregulation of pathways involved in cytosolic DNA-sensing, Toll-like receptor signaling, TNF signaling, and NF-kappa B signaling, all of which are downregulated by PG3@MSNs treatment (Supplementary Fig. 23). Furthermore, PG3@MSNs reverse the CpG DNA-disturbed lipid metabolism, downregulating lipid and atherosclerosis pathways while upregulating cholesterol metabolism (Supplementary Fig. 24), with gene set variation analysis (GSVA) further suggesting the involvement of proinflammatory response and lipid metabolism pathways in oxLOL-CpG DNA challenge and PG3@MSNs treatment (Fig. 5i and Supplementary Fig. 25).

Treated macrophages are classified into three subclusters based on signature genes using single-cell RNA sequencing analysis (Fig. 6a and Supplementary Fig. 26). Macro-1 represents an unchallenged cluster of primary macrophages, with the highest proportion in both the Control and oxLDL groups (Fig. 6b). CpG DNA stimulation under oxLDL conditions induces the emergence of Macro-2 and Macro-3, with Macro-2 becoming the predominant subcluster: The upregulation of *Tnf* and *Cxcl2* in Macro-2 suggests an activated, proinflammatory phenotype; In contrast, Macro-3 exhibits foamy signature, characterized by high expression of *Fabp4* and *Spp1*, which are involved in cholesterol ester formation and foam cell development in macrophages (Fig. 6b-d, and Supplementary Fig. 27) [33, 34]. After PG3@MSNs treatment, the proportions of both Macro-2 and Macro-3 are reduced (Fig. 6b). Monocle pseudotime analysis reveals macrophage progression from Macro-1 to Macro-2, culminating in Macro-3, with persistent upregulation of *Fabp4* and *Spp1* across pseudotime, suggesting these markers drive cfDNA-induced macrophage inflammatory-foamy differentiation (Fig. 6e, f and Supplementary Fig. 28). Immunofluorescence staining of aortic tissue from comorbidity models, using either systemic or local administration of PG3@MSNs, confirms the *in vitro* findings, showing that ligature-induced periodontitis elevates the expression of pro-inflammatory markers (*Tnf* and *Cxcl2*) and foamy markers (*Fabp4* and *Spp1*), while PG3@MSNs reduce their expression (Fig. 6g-j).

**Fig. 6.**
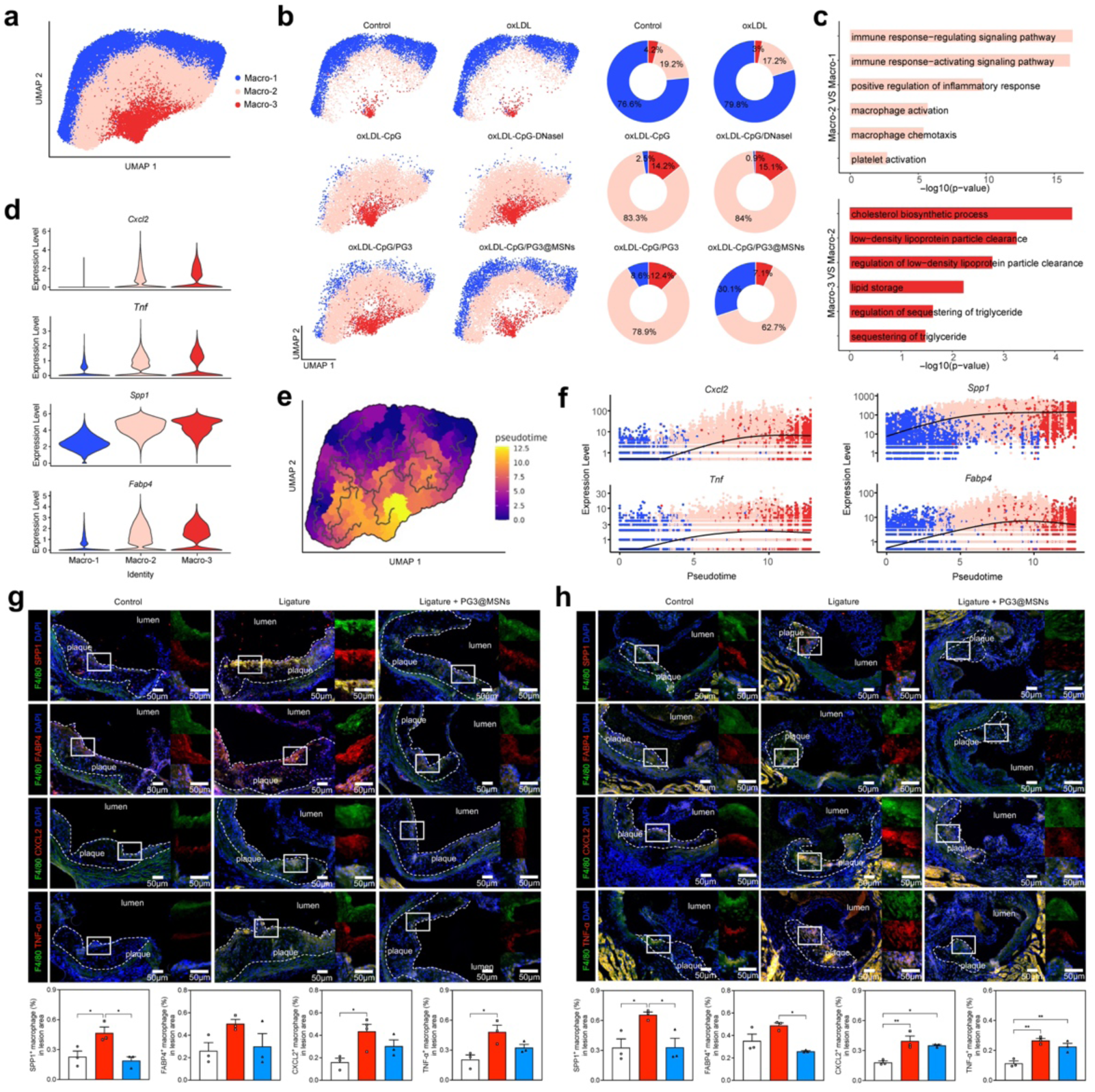
PG3@MSNs application helps prevent inflammatory foam cell formation in macrophages. a-b Uniform manifold approximation and projection (UMAP) of macrophages in the integrated group (**a**) and in control, LDL, oxLDL-CpG, oxLDL-CpG/DNase I, oxLDL-CpG/PG3, and oxLDL-CpG/PG3@MSNs groups (**b,** left), respectively. Changes in the proportion of macrophage subclusters in control, LDL, oxLDL-CpG, oxLDL-CpG/DNase I, oxLDL-CpG/PG3, and oxLDL-CpG/PG3@MSNs groups (**b,** right). **c** GO pathways enriched in Macro-2 (top) and Macro-3 (bottom). **d** Expression levels of inflammation- and foam cell-related genes *Cxcl2*, *Tnf*, *Spp1*, and *Fabp4* in different macrophage subclusters. **e** Monocle analysis of macrophages. **f** Expression levels of inflammation- and foam cell-related genes *Cxcl2*, *Tnf*, *Spp1*, and *Fabp4* in different macrophage subclusters along the differentiation trajectory in pseudotime. **g** Representative images of immunofluorescent staining of F4/80 (green) and SPP1, FABP4, CXCL2, or TNF-α (red) of the aortic valve in ApoE^-/-^ mice with systemic administration of PBS or PG3@MSNs in Fig 3 (top). Scale bars, 50 μm. Bottom, percentage of SPP1^+^/F4/80^+^, FABP4^+^/F4/80^+^, CXCL2^+^/F4/80^+^, or TNF^+^/F4/80^+^ in F4/80^+^ macrophages in the plaque lesion area (n = 3). **h** Representative images of immunofluorescent staining of F4/80 (green) and SPP1, FABP4, CXCL2, or TNF-α (red) of the aortic valve in ApoE^-/-^ mice with local administration of PBS or PG3@MSNs in Fig 4 (top). Scale bars, 50 μm. Bottom, percentage of SPP1^+^/F4/80^+^, FABP4^+^/F4/80^+^, CXCL2^+^/F4/80^+^, or TNF^+^/F4/80^+^ in F4/80^+^ macrophages in the plaque lesion area (n = 3). Data in **g** and **h** are means ± SEM; statistical analysis was performed using one-way ANOVA with Tukey’s post hoc test (**g**, **h**); \**P* < 0.05, \*\**P* < 0.01.

**Fig. 7.**
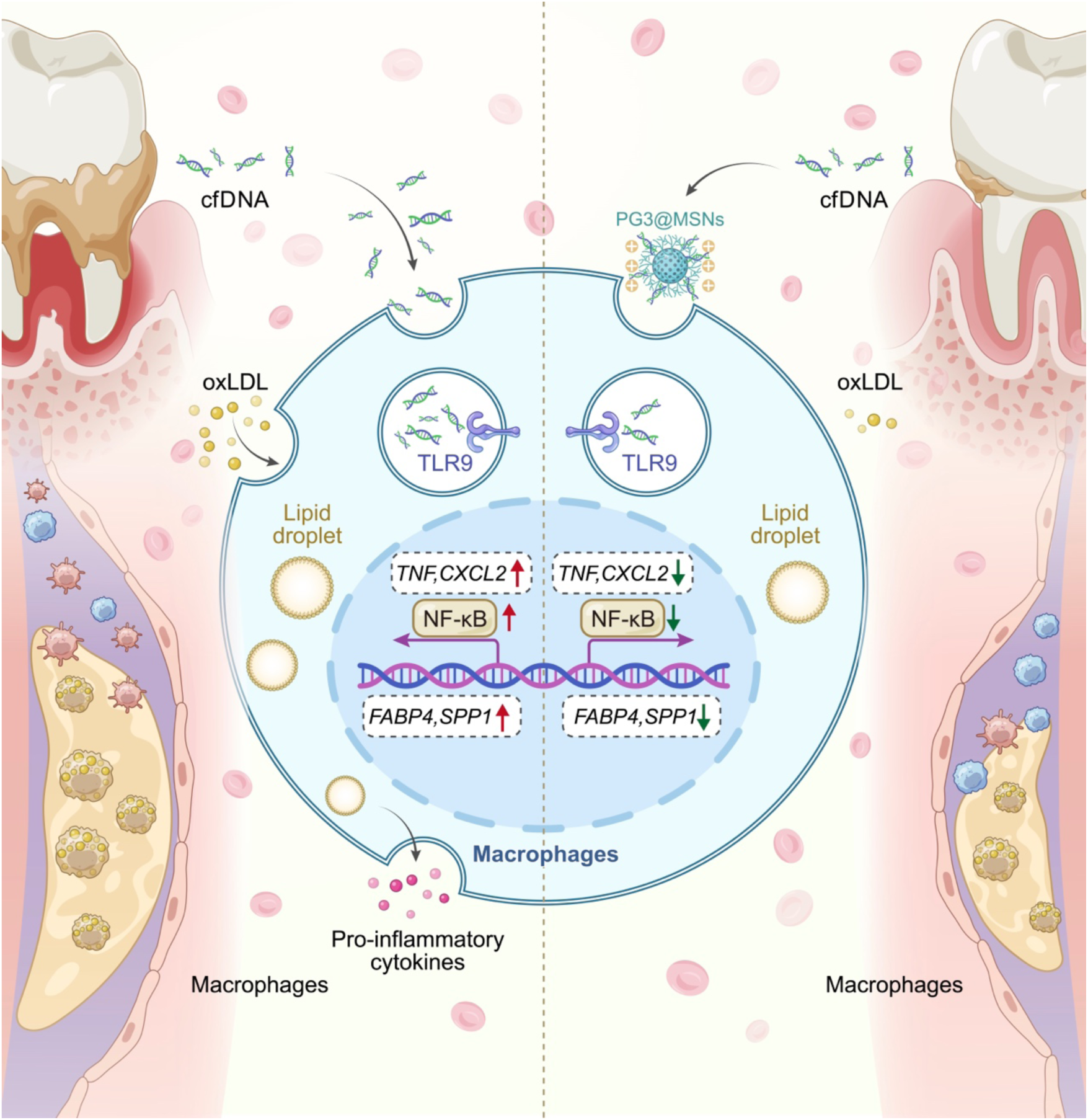
Schematic of cfDNA-based crosstalk between periodontitis and atherosclerosis. Periodontium-originated DNA plays a major role in atherosclerotic leision and cfDNA-capturing nanomaterials alleviates both periodontitis and atherosclerosis through cfDNA-TLR9 pathway and regulating inflammatory foam cell transformation in macrophages by modulating the expression of lipid-related foamy markers *Spp1* and *Fabp4*.

## Discussion

Understanding and controlling local-systemic disease crosstalk enables earlier detection, comprehensive treatment, disease prevention, and personalized healthcare, ultimately improving patient outcomes [35]. Periodontitis, as a typical local inflammation, stands out as a critical contributor [36]. Periodontitis can cause low-grade systemic inflammation [37, 38], which may contribute to the local-systemic disease connection. Control of periodontitis attenuates systemic inflammatory markers [39–43]. The relationship between periodontitis and linked comorbidities is often bidirectional, as systemic diseases can promote susceptibility to periodontitis [44]. Atherosclerosis, an essential systemic disease, is characterized by the accumulation of plaques within arterial walls, leading to the hardening and narrowing of arteries [45]. This process is driven by a combination of lipid metabolism dysregulation and chronic inflammation [46]. The resulting plaque buildup can restrict blood flow, significantly increasing the risk of cardiovascular events such as heart attacks and strokes [47]. Proinflammatory immune response play a crucial role in atherosclerosis [48], and emerging evidence suggests that local inflammatory conditions, like periodontitis, may exacerbate this systemic disease like atherosclerosis [49, 50].

To uncover the underlying mechanisms and develop mechanism-targeted treatment strategies, we focus on investigating the pathogenic factors involved in this local-systemic disease crosstalk and exploring innovative therapeutic approaches to break the cycle of periodontitis and atherosclerosis. Abnormally high levels of pathogenic cfDNA are recognized as a key factor in the pathogenesis of inflammatory diseases [51]. Our previous studies have demonstrated that DNA levels in circulation correlate with the severity of periodontal inflammation and controlling periodontal inflammation can help reduce these levels [52]. Similarly, elevated cfDNA has been documented in cardiovascular diseases [53]. In this study, we first examine the plasma cfDNA levels in patients with both periodontitis and atherosclerosis to explore the potential link between these local-systemic inflammatory comorbidities.

We strictly control the inclusion criteria for patient enrollment, and patients with any systemic disease other than atherosclerosis that may affect circulating cfDNA levels (such as diabetes, tumors, etc.) are excluded [54]. For patients with atherosclerosis, we use the angiography to confirm the diagnosis [55], and once atherosclerosis was confirmed, we conduct periodontal examinations. Interestingly, all patients diagnosed with atherosclerosis also exhibit periodontal inflammation. This contrasts with a previous report on DNA levels in patients with cardiovascular disease and periodontitis, likely due to differences in inclusion criteria [20]. As a chronic inflammatory condition, atherosclerosis may influence local inflammatory responses. Our findings demonstrate that plasma cfDNA levels consistently correlate with the severity of periodontitis, regardless of atherosclerosis presence; however, once atherosclerosis develops, plasma cfDNA levels do not correlate with the severity of atherosclerosis. These data again confirm that local inflammation can induce low-grade systemic inflammation, and suggest that timely control of local inflammation may help prevent the development of severe systemic inflammation; Once severe systemic inflammation is established, it can trigger a high systemic release of cfDNA, which becomes difficult to control and can significantly affect local inflammation.

Therefore, targeting cfDNA with functional nanomaterials may offer a promising approach to treating periodontitis-associated atherosclerosis by modulating this local-systemic disease crosstalk. Biodegradable mesoporous silica nanoparticles (MSNs) functionalized with cationic polymers are a promising option, as they have demonstrated the ability to capture DNA in the circulatory system and regulate inflammation [25]. Compared to the former report about using polyethylenimine as the functional group of the nanoparticle, PG3 is more promising with its 3D branched architecture, precise molecular weight, accessible internal cavities, and easy surface functionalization [56]. Here, we use third-generation polyamidoamine dendrimer (PG3)-functionalized MSNs (PG3@MSNs) to capture cfDNA. We conduct an *in vivo* study to validate the hypothesis that cfDNA originates from periodontal tissue and can be captured in systemic circulation. After locally injecting CpG DNA into periodontal tissue, we observe its presence in various organs, confirming its entry into the circulation. Following systemic administration of PG3@MSNs via intraperitoneal injection, we find that PG3@MSNs co-localize with CpG DNA in these organs, demonstrating the nanomaterial’s ability to capture periodontitis-driven cfDNA and its potential for targeted therapy in periodontitis-associated atherosclerosis.

Previous studies suggest that systemic administration to regulate the inflammatory response process triggered by periodontitis is a potential approach for alleviating atherosclerosis, as intravenous injection of tanshinone IIA and subcutaneous injection of the hTERT peptide fragment ameliorates the exacerbation of atherosclerosis in mice with periodontitis [57, 58]. Therefore, to confirm the efficacy of the cfDNA-targeting strategy in treating periodontitis-associated atherosclerosis, we established a murine model using Apoe^-/-^ mice with a western diet-induced atherosclerosis and ligature-induced periodontitis. Initially, we hypothesized that intraperitoneal injection of PG3@MSNs would only alleviate atherosclerosis but have no effect on local periodontitis, based on our previous study showing that cationic nanomaterials administered systemically for two weeks do not impact periodontitis [27]. However, after two months of systemic PG3@MSNs administration, we observed improvements in both atherosclerosis and periodontitis. This unexpected result may be due to the prolonged nature of the periodontitis-atherosclerosis model, suggesting that systemic inflammation also plays a role in local inflammation.

This finding prompts us to investigate the crosstalk between local periodontitis and systemic atherosclerosis, confirming that periodontitis aggravates atherosclerosis and that controlling periodontitis can help alleviate it. Evidence has demonstrated that active intervention in local periodontal factors is beneficial for the control of systemic vascular inflammation [59]. Thus, to explore this interaction, we administered local injections of PG3@MSNs into periodontal tissue to capture local cfDNA, maintaining the same dosing regimen in a murine model of atherosclerosis and periodontitis. Following local administration, plasma cfDNA levels decrease, and periodontal tissue destruction, alveolar bone loss, and vascular plaque formation are significantly inhibited. Next, we assess whether capturing systemic cfDNA in established atherosclerosis can inhibit local periodontal inflammation and alveolar bone loss. The results reveal that this approach does not significantly alleviate either local or systemic inflammation. This reinforces the idea from our clinical data that local inflammation drives systemic inflammation, emphasizing the importance of early control of local inflammation to prevent systemic inflammatory comorbidities. However, once systemic inflammation is established, systemic interventions appear less effective for both local and systemic inflammation. This aligns with the consensus report of periodontitis and cardiovascular diseases conducted by the European Federation of Periodontology and the World Heart Federation, highlighting that while preserving periodontal health can prevent cardiovascular diseases, its impact may be limited once such diseases have developed[60]. Involvement of immune cells in the atherosclerotic plaques has been uncovered by singe-cell landscape [61]. The addition of CpG DNA exacerbated the foamy change of oxLDL-treated macrophages, causing significant disturbances in immune response and lipid metabolism, suggesting an intimate relationship between lipids and inflammation [46]. Our findings also implied a foamy-like transformation from inflammatory macrophages, which aligns with the current perspective on atherosclerosis treatment, focusing on the resolution of inflammation [46, 62]. Through inhibition of inflammation activated by the cfDNA-TLR9-TNF signaling pathway, PG3@MSNs application helps prevent inflammatory foam cell formation in macrophages. Between the pro-inflammatory Macro-2 and inflammatory-foamy Macro-3 cells, the upregulation of *Spp1* and *Fabp4* genes was most prominent. Furthermore, *in vivo* results demonstrated that periodontitis significantly exacerbated the accumulation of SPP1^+^FABP4^+^ macrophages within arterial plaques, while treatment with PG3@MSNs successfully mitigated this effect.

Secreted Phosphoprotein 1 (SPP1), also known as Osteopontin (OPN), has been extensively reported to be associated with the progression of atherosclerosis [63]. Inflammatory cytokines, such as TNF-α, enhance the release of osteopontin (OPN) and further recruit immune cells, exacerbating local vascular inflammation. Fatty acid-binding protein 4 (FABP4) also plays a critical role in macrophage lipid metabolism by promoting the accumulation of cholesterol esters and inhibiting cholesterol efflux, significantly accelerating foam cell formation and thereby aggravating atherosclerosis [34]. Interestingly, we have reported in a muscle fibrosis model that *Spp1^+^*macrophages participate in inflammation-mediated muscle fibrosis by influencing *Fabp4^+^* fibro-adipogenic progenitor cells through paracrine signaling, and regulating inflammation by cfDNA-capturing nanoparticles can lead to an outstanding phenotypic transformation of both cell types, thereby promoting muscle regeneration [64]. Despite existing attempts to target SPP1 and FABP4 for cardiovascular disease treatment, satisfactory clinical outcomes have been lacking [65]. Our study demonstrates that the cationic nanomaterial PG3@MSNs mitigates inflammation by capturing cfDNA, thereby inhibiting the TLR9 pathway. This leads to the downregulation of SPP1 and FABP4 expression in macrophages, improving foam cell formation and reducing atherosclerotic lesions in mice. These findings provide evidence for the role of cfDNA as a mediator linking local and systemic inflammation and propose a novel strategy for the treatment of local-systemic inflammatory comorbidities.

There are several limitations of this study. To explore the impact of atherosclerosis on periodontitis, we conducted an animal experiment by inducing advanced atherosclerosis through a high-fat diet for three months, followed by periodontitis induction, which revealed more severe periodontal bone destruction in atherosclerotic mice compared to wild-type mice; however, the strength of our conclusions is limited due to confounding factors such as genetic deficiencies, dietary variations, and age-related periodontal changes. Future scRNA sequencing analysis of aortic tissue from mice with periodontitis-associated atherosclerosis is necessary to enhance our understanding of the underlying pathomechanisms and support the our findings. Although our experimental results indicate that cfDNA stimulation exacerbates lipid metabolism disorders in macrophages and promotes inflammatory responses, the role of specific cells and specific targets linking these effects still need gain- and loss-of-function studies to validate.

## Methods

### Materials and reagents

Polyamidoamine generation 3 (PAMAM-G3) (412422), ammonium nitrate (1.01187), tetraethyl orthosilicate (8.00658), (3-glycidyloxypropyl)trimethoxysilane (440167), bis[3-(triethoxysilyl)propyl]diselenide (15200), triethanolamine (8.22341), and cetyltrimethylammonium tosylate (CTAT) (8.14692) were purchased from Sigma-Aldrich (St. Louis, MO, U.S.A.). Dulbecco’s modified Eagle’s medium (DMEM) (11995065), Penicillin-Streptomycin (P/S) (10,000 U/mL) (15140122) were purchased from Gibco (Carlsbad, CA, U.S.A.). Fetal bovine serum (FBS) (A5669701) was purchased from GIBCO BRL (Grand Island, NY, USA). The CpG oligodeoxynucleotides (ODNs), including CpG 2006 (tlrl-2006) and CpG 1826 (either FITC-labeled or unlabeled, tlrl-1826f and tlrl-1826), and QUANTI-BlueTM solution (rep-qbs) were obtained from InvivoGen (San Diego, CA, U.S.A). Tribromoethanol (T903147) and tert-Amyl alcohol (A800283) were purchased from MAKLIN (Shanghai, China). H&E staining kit (G1120), Masson’s trichrome staining kit (G1346), Oil Red O staining kit (G1262), Total Cholesterol(TC) Content Assay Kit (BC1980) and Free Cholestenone(FC) Content Assay Kit (BC1890) were purchased from Solarbio (Beijing, China). Paraformaldehyde (4%) (BL539A) was purchased from Biosharp (Hefei, China). The Radio-Immunoprecipitation (RIPA) Lysis Buffer (K1020)was supplied by APExBIO (Houston, U.S.A). Cell Counting Kit-8 (C0028), BCA Protein Assay Kit (P0012), blocking buffer (P0260) and Antifade Mounting Medium with DAPI (P0131) were purchased from Beyotime (Shanghai, China). Quant-iT PicoGreen dsDNA assay kit (P11495), TRIzol (15596018) and anti-F4/80 Monoclonal Antibody (14-4801-85), and ELISA kit for mouse TNF-α and IL-6 (BMS607-3, BMS603-2) were purchased from Thermo Scientific (Waltham, Massachusetts, U.S.A). PrimeScript^TM^ FAST RT reagent Kit with gDNA Eraser (RR092A) and TB Green^®^ Premix Ex Taq™ (Tli RNaseH Plus) (RR420A) were purchased from Takara Bio (Beijing, China). Anti-TLR9 (BC05301367) was purchased from Bioss (Beijing, China). The anti-rabbit HRP-DAB IHC Detection Kit (SAP-9100) and DAB Color Development Kit (ZLI-9018) were purchased from ZSGO-BIO (Beijing, China). Oxidized-LDL(ox-LDL) (YB-002) was purchased from Yiyuan Biotechnology (Guangzhou, China). Anti-SPP1 antibody (A499), anti-FABP4 antibody (A25792), and anti-TNF-α antibody (A11534) were purchased from ABclonal Technology (Wuhan, China). FITC-conjugated Goat Anti-Rat IgG (SA00003-11) was purchased from Proteintech Group (Chicago, USA). Cy3-conjugated Goat Anti-Rabbit IgG (S0011) was obtained from Affinity Biosciences (Cincinnati, OH, U.S.A).

### Patients sample collection

130 consecutive subjects were evaluated for eligibility at West China Hospital of Stomatology and West China Fourth Hospital, Sichuan University, between November 2023 and September 2024. Based on the diagnostic criteria [66], they were classified as periodontal healthy individuals, gingivitis patients, or patients with periodontitis by oral examinations conducted by a trained periodontist. Periodontal clinical parameters recorded during the examination included PLI, BI and PD of the Ramfjord index teeth [67]. Atherosclerosis was diagnosed through coronary angiography, with the number and percentage of arterial stenosis recorded. Participants meeting any of the following criteria were excluded from the study: 1) pregnant or breastfeeding women; 2) individuals who have received localized radiation therapy to the oral cavity (such as head and neck radiotherapy); 3) those diagnosed with other systematic diseases in addition to coronary heart disease (CHD), such as diabetes, rheumatic diseases, etc.; 4) individuals with severe dysfunction of vital organs, including the heart, liver, kidneys, or lungs; 5) patients with malignant tumors; and 6) individuals unable to provide informed consent. The research followed the declaration of Helsinki on medical research guidelines reviewed in 2016. The study protocol was registered on the Chinese Clinical Trial Registry (ChiCTR2300077652), and ethical approval was obtained from the Ethics Committee of Sichuan University (WCHSIRB-CT-2019-12).

### Synthesis of diselenide-bridged MSN and PG3@MSNs

Diselenide-bridged MSNs were prepared as our protocol reported previously [68]. After dissolving 1.2 g of cetyltrimethylammonium tosylate (CTAT) and 0.3 g of triethanolamine in 80 mL of deionized water and stirring for an hour at 80°C. Then, 2.0 g of bis[3-(triethoxysilyl)propyl]diselenide (BTESePD) and 8.0 g of tetraethyl orthosilicate mixed with 3 mL of ethanol, and the mixture were added dropwise to the surfactant solution, which was then stirred at 80°C for 4 hours at 1000 rpm. Following centrifugation, the products were collected, and the reaction mixture was washed multiple times with ethanol and refluxed in a solution of ethanol of ammonium nitrate (1% w/v) for 12 hours to remove the surfactant CTAT. The as-synthesized diselenide-bridged MSN were gathered, rinsed, and dried for subsequent experiments.

For synthesizing PG3@MSNs, 1.0 mg of MSNs was dispersed in toluene (250 mL), followed by the addition of (3-glycidyloxypropyl)trimethoxysilane (1.5 mL). The mixture was then refluxed at 80 °C for 24 h to yield epoxysilane-functionalized MSN. Following purification, the epoxysilane-functionalized MSN (500 mg) were dispersed in PAMAM-G3 solution (250 mL, 1 mg mL^−1^) and stirred at room temperature for 24 h. The products were collected by centrifugation at 10000 rpm for 15 minutes, washed three times with water to obtain PG3@MSNs.

### Characterization of MSNs and PG3@MSNs

The structural features of MSNs and PG3@MSNs were visualized via transmission electron microscopy (TEM, JEOL, Ltd., Japan) and scanning electron microscopy (SEM, FEI Quanta 200F). The particle size and size distribution of PG3@MSNs were measured using a particle size analyzer based on the dynamic light scattering (DLS) principle. The surface zeta potential was determined using electrophoretic light scattering. The specific surface area of PG3@MSNs was determined using a surface area and porosity analyzer based on the Brunauer-Emmett-Teller (BET) method. The PG3@MSNs content within PG3@MSNs was quantified by thermogravimetric analysis (TGA, PerkinElmer, U.S.A.).

### DNA binding assay

The binding ability of MSNs, PG3, and PG3@MSNs with ct-DNA was evaluated as described previously [68]. ct-DNA solution, PicoGreen, and MilliQ water were mixed in a 96-well plate. Followed by shaking and adding different concentrations of MSNs, PG3, or PG3@MSNs solutions. After 1-hour incubation at 37℃, the fluorescence intensity was measured with a multi-mode microplate reader (SpectraMax iD3, Molecular Devices).

### Cytotoxicity assay *in vitro*

HEK-Blue™ TLR9 reporter cells were seeded at a density of 8 × 10^4^ cells per well, and RAW 264.7 cells were seeded at 2 × 10^4^ cells per well in 96-well culture plates. The cells were cultured until full adhesion was achieved. Subsequently, the cells were treated with PG3, MSNs, and PG3@MSNs at concentrations of 1 μg/ml, 10 μg/ml, 20 μg/ml, 40 μg/ml, 80 μg/ml, 120 μg/ml, 240 μg/ml, 480 μg/ml, and 960 μg/ml for 24 hours. Cell viability was then assessed according to the manufacturer’s instructions for the CCK-8 assay kit.

### *In vitro* anti-TLR9 activation and anti-inflammation assay

HEK-Blue™ TLR9 reporter cells were seeded at a density of 8 × 10^4^ cells per well and RAW 264.7 cells were seeded at a density of 2 × 10^4^ cells per well in a 96-well culture plate and incubated with basal DMEM for 12 hours. In the materials-treated groups, MSNs, PG3, or PG3@MSNs (10 μg/mL) were introduced 30 minutes before the CpG 2006 (for HEK-Blue™ TLR9 reporter cells, 1 μg/mL) or CpG 1826 (for RAW 264.7 cells, 1 μg/mL) agonist. Following a 24-hour incubation period, supernatants of HEK-Blue™ TLR9 reporter cells were collected, and the activity of embryonic alkaline phosphatase (SEAP) was measured with the QUANTI-Blue assay kit; supernatants of RAW 264.7 cells were collected and levels of TNF-α and IL-6 in the supernatants were quantified.

For *in vitro* anti-TLR9 activation and anti-inflammatory experiments utilizing samples from *in vivo* studies, HEK-Blue™ TLR9 reporter cells and RAW264.7 cells were treated with PG3@MSN (10 μg/ml) in a volume of 200 μl for 30 minutes. Subsequently, 5 μl of plasma from 17-week-old C57BL/6 mice (C57), plasma from ApoE^-/-^ mice in the Control group (ApoE^-/-^), and plasma from ApoE^-/-^ mice in the Ligature group (PD/ApoE^-/-^) were added for 24-hour stimulation, respectively. The activation of HEK-Blue™ TLR9 reporter cells and the secretion of TNF-α and IL-6 by RAW 264.7 cells were assayed as previously described.

### Mice

Balb/c mice (male, eight weeks old), ApoE^-/-^ mice (male, eight weeks old), and C57BL/6 mice (male, eight weeks old) were purchased from Dossy Experimental Animals Co., Ltd. (Chengdu, Sichuan, China) and were randomly divided into groups for interventions according to the study designs. The animal studies were approved by the Ethical Committee of the West China Hospital of Stomatology, Sichuan University (NO. WCHSIRB-D-2019-122).

### Biodistribution of locally injected CpG DNA and intraperitoneally injected PG3@MSNs

Eight-week-old male Balb/c mice were divided into control and material-treated groups. The material-treated group received a concurrent injection of FITC-labeled CpG DNA (100 μg/ml) into the periodontal tissues at six sites (5 μL/site) accomplished using a microsyringe (25 μL, 32-gauge needle) under anesthesia with isoflurane, and an intraperitoneal injection of Cy7-labeled PG3@MSNs (1 mg/ml, 10 mg/kg) [25]; while the control group was injected with an equal volume of saline. At 2 hours, 12 hours, and 24 hours post-injection, both the anesthetized mice and their major organs (maxilla, heart, liver, spleen, lung, and kidney) were carried with small animal in vivo imaging system (IVIS) (excitation wavelength: 710 nm; emission wavelength: 780 nm) to detect the biodistribution of locally injected CpG DNA and intraperitoneally injected PG3@MSNs. The other half of the mice were euthanized for major organs (heart, liver, spleen, lung, and kidney) collection. The organs were fixed in 4% paraformaldehyde at 4°C under light-protected conditions, followed by OCT embedding and frozen sectioning. These samples were subsequently used for immunofluorescence staining and observation.

### In vivo studies for comorbidities treated with PG3@MSNs

Eight-week-old male ApoE^-/-^ mice were chosen. All mice were using the Western diet to induce atherosclerosis. For the mice with periodontitis, a 5-0 suture was inserted and ligatured around the cervix of the maxillary second molar with general anesthesia by sterile avertin (tribromoethanol: 200 mg/10 mL kg^−1^, dissolved in deionized water). The ligatures were checked during each administration, and any loose or detached sutures were promptly replaced. Maxilla, the entire thoracoabdominal aorta, heart, and other main organs were collected after the experiments following different designs.

For the comorbidity mice model with systemic administration of nanomaterials, ApoE^-/-^ mice were divided into four groups: group without ligature (Control), untreated group with ligature and PBS injection (Ligature), PG3@MSNs treated group with ligature and PG3@MSNs injection (Ligature+PG3@MSNs), and PG3@MSNs treated group without ligature (PG3@MSNs). All mice received intraperitoneal injections of PBS or PG3@MSNs on alternate days after ligature completion. The concentration of PG3@MSNs was referenced from the previous report (1 mg/ml, 10 mg/kg) [25]. Mice were sacrificed after eight weeks, and samples were collected for further analysis.

For the comorbidity mice model with local administration of nanomaterials, ApoE^-/-^ mice were divided into three groups: group without ligature (Control), untreated group with ligature and PBS injection (Ligature), and PG3@MSNs treated group with ligature and PG3@MSNs injection (Ligature+PG3@MSNs). Local administration of PBS or PG3@MSNs (1 mg/mL, 10 mg/kg) was accomplished using a microsyringe (25 μL, 32-gauge needle) into maxillary gingiva at six sites (5 μL/site) around the ligatured second molar under anesthesia with isoflurane [27]. To minimize soft tissue damage, periodontal microinjections were given every three days, with PBS and PG3@MSNs noninvasively applied around the ligatured molar on non-injection days. Mice were sacrificed after four weeks, and samples were collected for further analysis.

To assess the preventive effect of PG3@MSNs on periodontitis in atherosclerotic mice, ApoE^-/-^ mice were fed a western diet for 3 months to induce severe atherosclerosis, followed by 2 weeks of systemic administration with PBS or PG3@MSNs. C57BL/6 mice with periodontitis but without atherosclerosis were also included for comparison. Both C57BL/6 mice and atherosclerotic ApoE^-/-^ mice were established with ligature-induced periodontitis (AS + Ligature group and AS + PG3@MSNs + Ligature group) and sacrificed for sample collection after two weeks.

### Sample collection

Each maxilla was surgically divided into two parts, one of which was fixed and decalcified in a Formalin-EDTA Decalcifying Solution at 4 °C for four weeks (changing the EDTA solution twice during this time). Then, the decalcified maxilla was dehydrated, embedded in paraffin wax, and sectioned into 4 μm thick slices for further histological staining. In addition, gingival tissue was carefully separated from the other side of the maxilla and stored at -80 °C till RNA and protein extraction. The rest maxillary bone was scanned with Micro-CT for further analysis.

The thoracoabdominal aortae used for lipid staining were fixed in 4% paraformaldehyde at 4 °C and subsequently stained with Oil Red O. The other aortae were bisected and stored at -80°C for subsequent DNA and protein extraction.

Heart samples were fixed in 4% paraformaldehyde and subsequently dehydrated in 30% sucrose solution at 4 °C for 24 hours for frozen sectioning. Two mm of the cardiac apex was trimmed perpendicular to the long axis of the heart, with the cut surface of the apex facing downwards. The ascending aortic segment was lifted, and the heart was uprightly embedded in the OCT compound on dry ice. The frozen OCT block was fixed on a cryostat for continuous sectioning at a thickness of 8 μm. Sections containing the aortic valve structure were collected on adhesive slides, labeled, and stored at - 20°C for further staining.

The collected major organs (kidney, liver, spleen, and lung) of mice with comorbidities were fixed in 4% paraformaldehyde, dehydrated, embedded in paraffin wax, and sectioned into 4 μm thick slices for further histological staining.

### Micro-CT reconstruction and bone resorption quantitative analyses

Fixed maxillary bone samples were scanned with micro-CT (145 mA, 55 kVp, 10 μm slice thickness) (vivaCT 80, SCANCO Medical, Switzerland). Three-dimensional reconstructions were performed using Mimics Research 19.0.0 (Materialise, Belgium). The bone resorption area (mm²) between the cemento-enamel junction (CEJ) and alveolar bone crest (ABC) was quantified using Image Pro Plus (Media Cybernetics, USA). Trabecular bone parameters in the furcation region of the maxillary second molars were analyzed with customized scripts integrated into the vivaCT 80 system.

### Vessel Oil Red O gross staining

Followed by trimming of the extravascular connective tissue, the vessel was then longitudinally opened and flattened using microscissors for direct subsequent Oil Red O gross staining following the manufacturer’s instructions.

### Histological analysis

After dewaxing and dehydrating, the paraffin sections of maxillae and major organs were stained for H&E using a staining kit. The frozen sections of atherosclerotic plaque in the aortic sinus were stained for H&E, Oil Red O, and Masson using kits following the manufacturer’s instructions. Tissue sections were imaged under an optical microscope at different objective magnifications. Image Pro Plus software (Media Cybernetics, USA) was used to calculate and analyze plaque size, lipid deposition area, and fiber content proportion in gross aortic staining and aortic valve sections.

### Immunohistochemistry analyses

The immunohistochemistry (IHC) analysis was performed to detect the expression and location of TLR9 in both periodontal tissue and aortic plaque. Briefly, a primary antibody of a rabbit polyclonal antibody to TLR9 (1:100) was applied to the sections overnight. Then, the sections were subjected to incubation with secondary antibodies, followed by development using the 3,3′-diaminobenzidine chromogens in accordance with the established methodology. Nuclei were visualized with hematoxylin. Images were photographed by slide scanner (SLIDEVIEW VS200, Olymbus, Japan) and quantitatively analyzed by ImageJ software. Relative quantitative assessments were then performed by comparing the integral optical density (IOD) of the TLR9 positive staining areas.

### Immunofluorescence analyses

For the detection of biodistribution of FITC-labled CpG and Cy5-labled PG3@MSNs, sections of major organs were mounted with DAPI containing mounting medium to visualize nuclei. The frozen sections of the heart at the aortic valve level were stained for macrophage marker F4/80, along with SPP1, FABP4, TNF-α, or CXCL2. After a 15-minute incubation at room temperature in QuickBlock™ Blocking Buffer, the sections were incubated overnight at 4 °C with the first primary antibody, anti-F4/80 (1:50), for macrophage staining. Subsequently, sections underwent 1-hour incubation at room temperature with the first fluorescent secondary antibody, FITC-conjuncted Goat Anti-Rat IgG (1:200). Subsequently, the sections were incubated for 1 hour at room temperature with the first fluorescent secondary antibody, FITC-conjugated Goat Anti-Rat IgG (1:200). A second round of antigen retrieval and blocking was performed, followed by overnight incubation at 4 °C with a second primary antibody: anti-F4/80 (1:50), anti-SPP1 (1:100), anti-FABP4 (1:200), anti-TNF-α (1:100), or anti-CXCL2 (1:100). The sections were then incubated with the second fluorescent secondary antibody, Cy3-conjugated Goat Anti-Rabbit IgG (1:200). Finally, the nuclei were visualized using DAPI. All stained sections were scanned with Olympus VS200 fluorescence microscope (Olympus, Japan) and quantitatively analyzed by ImageJ software.

### RNA and protein extraction from animal samples

Samples were homogenized utilizing a high-speed tissue grinder (KZ-II; Servicebio, Wuhan, Hubei, China). Total RNA was extracted employing the RNApure Total RNA Fast Extraction Kit, and its quality and concentration were evaluated using a Nanodrop 2000 spectrophotometer (Thermo Scientific, Waltham, Massachusetts, USA). Subsequently, reverse transcription was conducted using the PrimeScript^®^RT reagent kit with gDNA eraser, with the resultant complementary DNA (cDNA) stored at −20°C for further testing. Total protein was extracted using PathScan Sandwich ELISA Lysis Buffer. The total protein concentration was determined by the bicinchoninic acid (BCA) method. The protein samples were then stored at −80°C for further testing.

### Quantitative real-time polymerase chain reaction assay

Total RNA was extracted using TRIzol reagent, and 1 μg of this RNA was reverse-transcribed using an PrimeScript^®^RT reagent Kit with gDNA Eraser. Quantitative polymerase chain reaction (PCR) was subsequently carried out utilizing TB Green® Premix Ex Taq^TM^ II Kit. The amplified transcripts were quantified employing the comparative Ct (threshold cycle) method.

### Cytokine concentration analysis

The concentrations of TNF-α and IL-6 in plasma, culture medium supernatants, and total tissue protein were determined using ELISA kits according to the manufacturer’s instructions.

### Plasma biochemical parameter analysis

The levels of total cholesterol (TC), total triglycerides (TG), and high-density lipoprotein cholesterol (HDL-c) were assayed using a chemistry analyzer (Chem-ray-800, Rayto), with reagents and settings recommended by the manufacturer.

### *In vitro* foam cell formation induction assay

RAW 264.7 cells were seeded at a density of 4 × 10^5^ cells per well in a 6-well culture plate and incubated with basal DMEM for 12 hours. Then, 50µg/mL of ox-LDL was utilized to induce the formation of foam cells in RAW 264.7 macrophages. In the materials-treated groups, DNase I, PG3, or PG3@MSNs (10 μg/mL) were introduced 30 minutes before the oxLDL and CpG 1826 (1 μg/mL) agonist for a 48-hour incubation period.

### Oil Red O staining and cellular cholesterol measurement of cells

Oil Red O staining was conducted to visualize lipid accumulation within macrophages. For the Oil Red O staining procedure, the cells were initially fixed with 4% PFA and then stained with Oil Red O. To quantify the intracellular levels of total cholesterol (TC) and free cholesterol (FC), the Cholesterol Quantitation Kit (Solarbio, China) was employed, following the manufacturer’s guidelines. The content of cholesteryl esters (CE) was calculated by subtracting the FC content from the TC content.

### Transcriptome sequencing

Total RNA was extracted from the RAW cell samples with TRIzol reagent according to the manufacturer’s protocol. Then, mRNA was enriched using magnetic beads coated with Oligo(dT). A fragmentation reagent was added to cleave the mRNA into short fragments. Using the fragmented mRNA as a template, first-strand cDNA was synthesized with hexamer random primers. Subsequently, a second-strand synthesis reaction was set up to generate double-stranded cDNA, which was purified using a kit. The purified double-stranded cDNA underwent end repair, A-tailing, and adaptor ligation, followed by size selection of the fragments. Finally, PCR amplification was performed. The constructed library was quality checked using an Agilent 2100 Bioanalyzer before sequencing on a sequencer, generating 125 bp or 150 bp paired-end data. The transcriptome sequencing was conducted by OE Biotech Co., Ltd. (Shanghai, China) and further bioinformatic analysis was performed with the help of OECloud tools at https://cloud.oebiotech.com.

### Single-cell RNA-seq data preprocessing and analysis

The MobiVision software pipeline (version 1.1) provided by MobiDrop was used to demultiplex cellular barcodes, map reads to the genome and transcriptome using the STAR aligner, and down-sample reads as required to generate normalized aggregate data across samples, producing a matrix of gene counts versus cells. We processed the unique molecular identifier (UMI) count matrix using the R package Seurat [69] (version 5.0.0). To remove low-quality cells and likely multiplet captures, which is a major concern in microdroplet-based experiments, a set of criteria were conducted: Cells were filtered by (1) gene numbers (gene numbers < 200), (2) UMI (UMI <1000), (3) log10GenesPerUMI (log10GenesPerUMI < 0.7), (4) percentage of mitochondrial RNA UMIs (proportion of UMIs mapped to mitochondrial genes > 10%) and (5) percentage of hemoglobin RNA UMIs (proportion of UMIs mapped to hemoglobin genes > 5%). Subsequently, DoubletFinder package (version 2.10-0) was applied to identify potential doublet [70]. After applying these QC criteria, single cells were included in downstream analyses. Individual samples were processed by log-normalization, the top 2,000 features were identified, the data was scaled, a principal component analysis (PCA) was computed, and the data was clustered via Louvain clustering using the top 30 principal components (PCs). To mitigate the effects of cell cycle heterogeneity on cell clustering, each sequenced cell was given a cell cycle phase ‘score’ based on the expression of canonical markers using the Seurat function ‘CellCycleScoring’, with the Seurat object as input in addition to G1/S- and G2/M-phase specific genes [71]. Differentially expressed genes in each cluster were identified using the FindAllMarkers function in Seurat v5. To remove the batch effects in single-cell RNA-sequencing data, the mutual nearest neighbors(MNN) was performed [72]. Cells were annotated by marker gene expression. The full dataset was processed as above. Differentially expressed genes (DEGs) were selected using the FindMarkers function (test.use = presto) in Seurat [73]. P adj < 0.05 and |log2foldchange| > 0.25 was set as the threshold for significantly differential expression. GO enrichment of DEGs were respectively performed using R-based package clusterProfiler (version 4.10.0) [74]. Significantly enriched pathways (adjusted p-value < 0.05) were selected to present in the Figure. Monocle 3 (version 1.0.0) was used to construct single-cell trajectories [75]. Cells from annotated clusters were extracted and reprocessed (including normalization and batch effect correction, dimensionality reduction and clustering) with Monocle 3. The trajectory was then learned. We selected the beginning nodes of the trajectory where more adjacent cells come from cluster Macro-1. Modules of interested genes were then calculated using plot_genes_in_pseudotime function. Nearly all graphical Figures were generated using R v4.4.1 and subsequently edited in Adobe Illustrator.

### Statistical analysis

Statistical analyses were conducted using GraphPad Prism 9.0 software (GraphPad, U.S.A). All values were expressed as mean ± standard error of the mean (SEM), with a minimum of three biological replicates. Assumptions of normality and homogeneity of variances were tested prior to conducting statistical analyses. For normally distributed data, comparisons between two groups were made using the Student’s t-test; while a oneway analysis of variance (ANOVA) followed by Tukey’s multiple comparisons test was used for comparisons among multiple groups. Correlations between clinical variables were assessed using Spearman’s correlation coefficient (r). A *P*-value < 0.05 was considered statistically significant. Statistical significance levels were indicated as follows: *P* < 0.05 (*), *P* < 0.01 (**), *P* < 0.001 (***), and *P* < 0.0001 (****).

