## Supplemental Table 1-2 and Supplemental Figure 1-28 for "Therapeutic targeting inflammation linking periodontitis and atherosclerotic comorbidities using cell-free DNA-capturing nanomaterials"

**Table S1. Basic information of subjects and periodontal clinical parameters**

| Characteristics | Groups | | | $p$ value (among three groups) | $p$ value (between PD & PD+AS) |
|---|---|---|---|---|---|
| | Healthy | Periodontitis (PD) | Periodontitis + Atherosclerosis (PD+AS) | | |
| Number of subjects ($n$) | 25 | 27 | 25 | | |
| Male / female | 16/9 | 11/16 | 15/10 | 0.2001 | 0.1717 |
| Age (years) | 25.60±1.61 (range 25-32) | 34.85±11.88 (range 25-60) | 56.04±6.66 (range 33-65) | <0.0001 | <0.0001 |
| Max PD (mm) | 2.92±0.28 | 6.89±2.01 | 6.00±1.53 | <0.0001 | 0.0801 |
| Mean PD (mm) | 1.87±0.31 | 3.42±0.67 | 3.66±0.70 | <0.0001 | 0.2226 |
| PLI | 0.57±0.29 | 1.78±0.46 | 1.60±0.65 | <0.0001 | 0.2698 |
| Plasma cfDNA (ng/ml) | 267.49±65.93 | 345.00±131.74 | 471.41±71.01 | <0.0001 | <0.0001 |
| NLR | 2.05±1.00 | 2.56±1.32 | 2.85±1.43 | 0.0791 | 0.4746 |
| PLR | 132.49±32.26 | 128.93±55.96 | 152.17±72.28 | 0.2876 | 0.2010 |
| LMR | 5.30±1.43 | 4.44±1.63 | 4.37±1.66 | 0.0732 | 0.8841 |
| SII | 509.06±231.88 | 583.58±435.21 | 556.82±352.70 | 0.9726 | 0.8852 |

PD, pocket probing depth; PLI, plaque index; NLR, neutrophil and lymphocyte ratio; PLR, palate and lymphocyte ratio; LMR, lymphocyte and monocyte ratio; SII, systematic inflammation index.

**Table S2. Correlation between plasma cfDNA concentration and periodontal clinical parameters**

| Statistics | Correlation Coefficient | | |
|---|---|---|---|
| | Mean PD (mm) | Max PD (mm) | PLI |
| $R^2$ Value | 0.2135 | 0.1025 | 0.06513 |
| *P* Value | <0.0001 | 0.0045 | 0.0251 |

PD, pocket probing depth; PLI, plaque index.

**Table S3. Sequences of the primers used for RT-qPCR**

| | |
|---|---|
| *Gapdh -F* | *AGGTTGTCTCCTGCGACTTCA* |
| *Gapdh -R* | *CCAGGAAATGAGCTTGACAAA* |
| *Tlr9-F* | *TTCTCAAGACGGTGGATCGC* |
| *Tlr9-R* | *GCAGAGGGTTGCTTCTCACG* |
| *Myd88-F* | *CATGGTGGTGGTTGTTTCTGAC* |
| *Myd88-R* | *TGGAGACAGGCTGAGTGCAA* |
| *Traf6-F* | *TGTTCTTAGCTGCTGGGGTGT* |
| *Traf6-R* | *GAAGGAGCTGGAGAGGTTCC* |
| *Rela-F* | *TTCCTGGCGAGAGAAGCAC* |
| *Rela-R* | *AAGCTATGGATACTGCGGTCT* |
| *Tnf-F* | *AGGGTCTGGGCCATAGAACT* |
| *Tnf-R* | *CCACCACGCTCTTCTGTCTAC* |
| *Il6-F* | *CTCTGCAAGAGACTTCCATCCAGT* |
| *Il6-R* | *GAAGTAGGGAAGGCCGTGG* |
| *Sr-a1-F* | *GCGGGAGGTCCTGTATGACT* |
| *Sr-a1-R* | *CGTCGAGACCCTTTCTCCCT* |
| *Acat1-F* | *TCCACTCCATGCACCACAGTAAAC* |
| *Acat1-R* | *CGCCTGCCACCATCACATCC* |
| *Abca1-F* | *AGAAGGAGGCTCGGCTGAAGG* |
| *Abca1-R* | *GAGGGATGAGGCTGCTAACAAACC* |

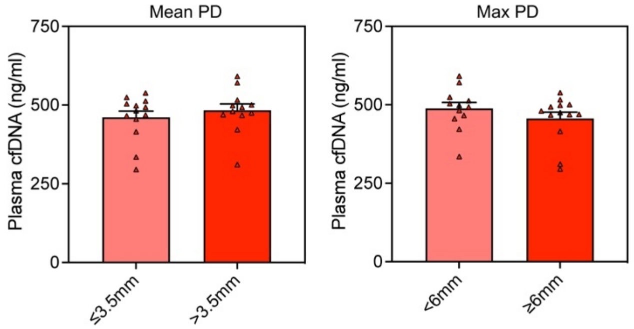


**Supplementary Fig. 1** Plasma cfDNA concentrations in subgroups of patients with periodontitis complicated by atherosclerosis, categorized by different mean PD and max PD. Statistical significance is calculated via a two-tailed Student's *t*-test.

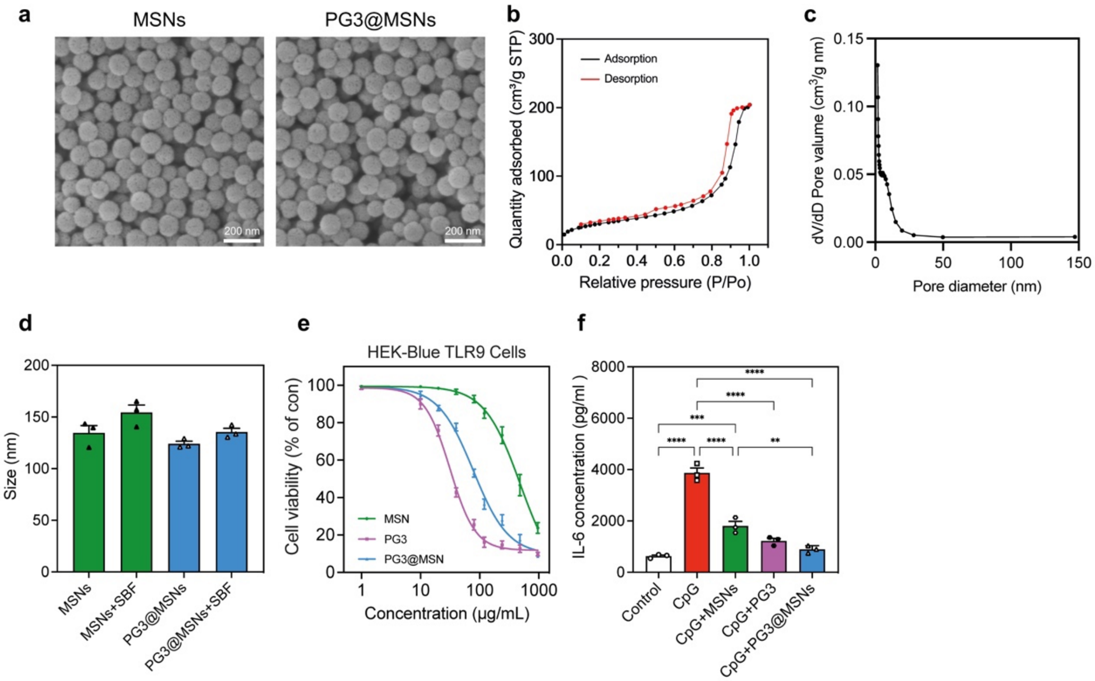


**Supplementary Fig. 2 The characterization and in vitro activity of PG3@MSNs.**

**a** SEM imaging of MSNs before and after coating with PG3. Scale bar, 200 nm.

**b** Size measurement of MSNs and PG3@MSNs with of without SBF (n = 3). SBF, simulated body fluids.

**c** BET analysis of SeHANs and G3@SeHANs.

**d** Viability of HEK-Blue TLR9 reporter cells treated for 24 hours with various concentrations of MSNs, PG3, and PG3@MSNs.

**e** RAW 264.7 macrophages were stimulated with CpG DNA (ODN 1826) in the absence or presence of MSNs, PG3, and PG3@MSNs for 24 hours. Supernatants were assayed for IL-6 by ELISA.

Data are means ± SEM; differences were assessed by one-way analysis of variance and Tukey's multiple comparisons test. *$P < 0.05$, **$P < 0.01$, ***$P < 0.001$, ****$P < 0.0001$.

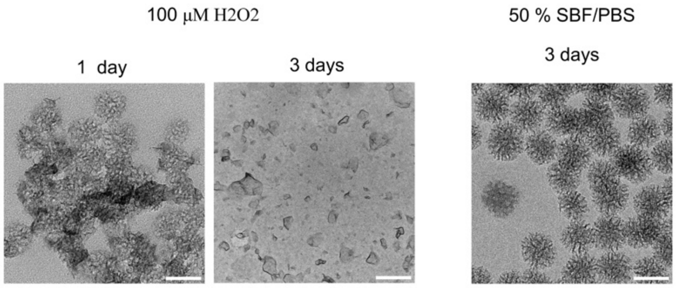


**Supplementary Fig. 3 Degradation of PG3@MSNs.**

Representative TEM images of PG3@MSNs after 1 day and 3 days incubation in $H_2O_2$ (left) and 3 days incubation in 50% SBF/PBS (right). Scale bar, 100 nm.

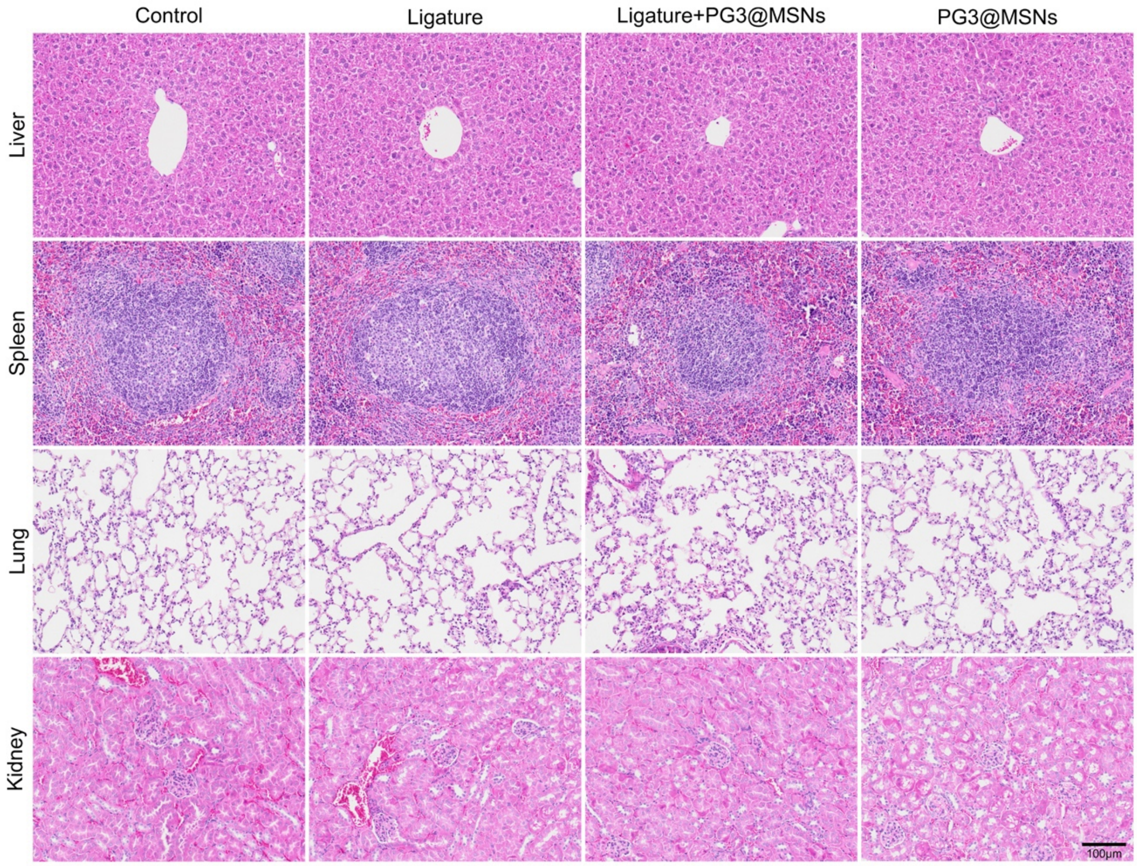


**Supplementary Fig. 4 H&E staining of major organs.**

Representative images of H&E staining of major organs of mice in **Fig.3**. Scale bar, 100 μm.

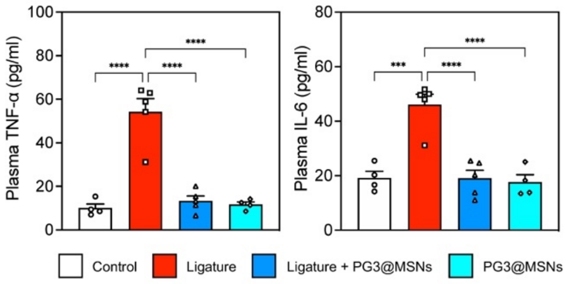


**Supplementary Fig. 5 Quantification of plasma TNF-α and IL-6 concentrations using ELISA.**
n = 4 in Control and PG3@MSNs, n = 5 in Ligature and Ligature + PG3@MSNs. Data are means ± SEM; differences were assessed by one-way analysis of variance and Tukey's multiple comparisons test. ***$P < 0.001$, ****$P < 0.0001$.

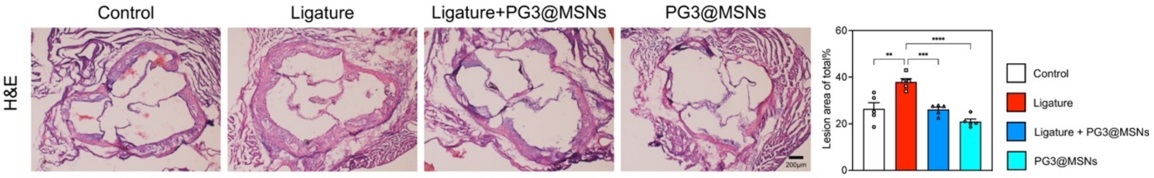


**Supplementary Fig. 6 H&E staining of aortic valve.**
**a** Representative images of H&E staining of aortic valve. Scale bar, 200 μm.
**b** Quantification of plaque occupying the lumen area in **a** (n = 5).
Data are means ± SEM; differences were assessed by one-way analysis of variance and Tukey's multiple comparisons test. **$P < 0.01$, ***$P < 0.001$, ****$P < 0.0001$.

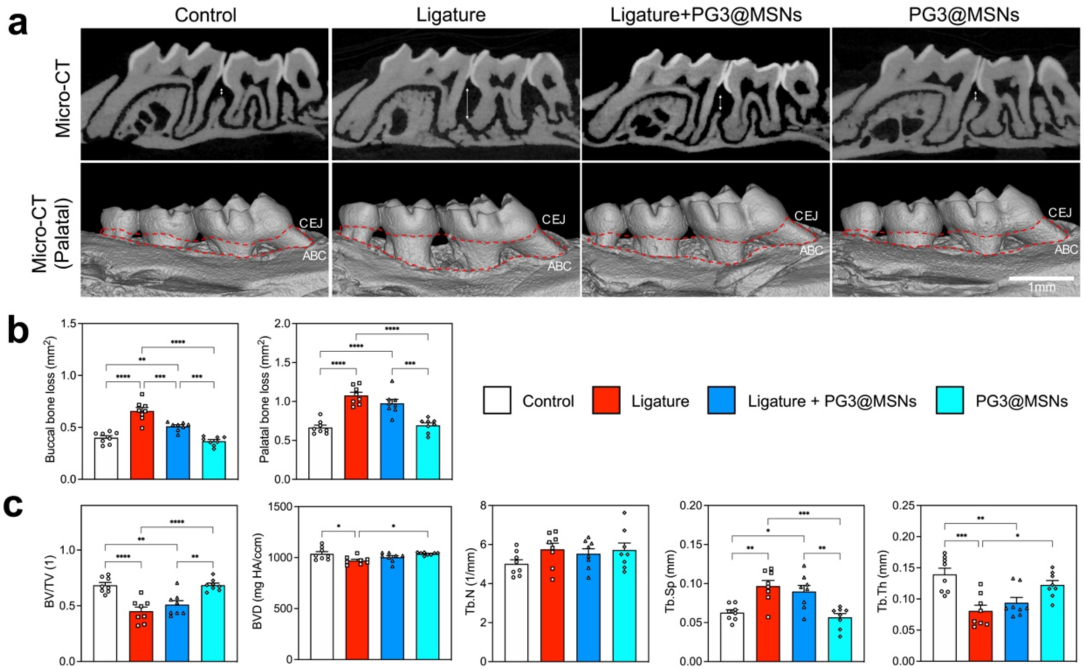

**Supplementary Fig. 7 Micro-CT of periodontal tissue.**

**a** Resliced view (top) and palatal view (bottom). Scale bars, 1 mm.

**b** Quantification of buccal and palatal alveolar bone resorption (n = 8).

**c** Quantification of trabecular bone parameters in the furcation area of the maxillary second molar. BV/TV, bone volume fraction. BVD, bone volume density. Tb.N, trabecular number. Tb.Sp, trabecular separation. Tb.Th, trabecular thickness.

Data are means ± SEM; differences were assessed by one-way analysis of variance and Tukey's multiple comparisons test. $^*P < 0.05$, $^{**}P < 0.01$, $^{***}P < 0.001$, $^{****}P < 0.0001$.

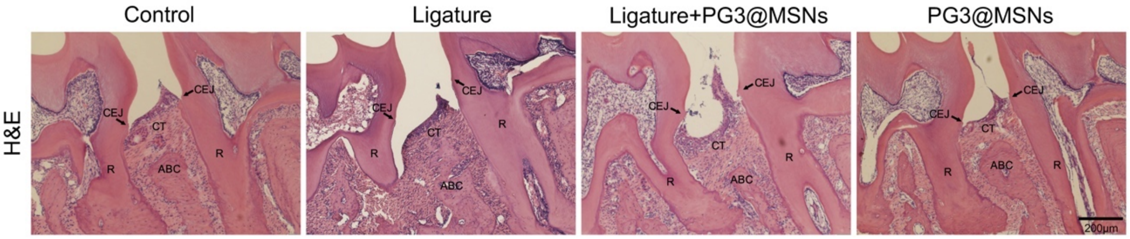


**Supplementary Fig. 8 H&E staining of periodontal tissue.** Scale bar, 200 μm.

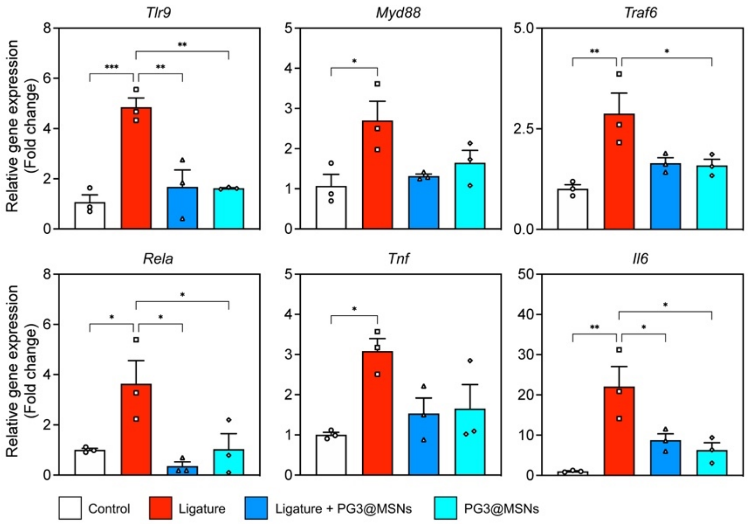


**Supplementary Fig. 9 Relative gene expression of *Tlr9*, *Myd88*, *Traf6*, *Rela*, *Tnf*, and *Il6* in thoracoabdominal aorta.**

n = 3, Data are means ± SEM; differences were assessed by one-way analysis of variance and Tukey's multiple comparisons test. $^*P < 0.05$, $^{**}P < 0.01$, $^{***}P < 0.001$.

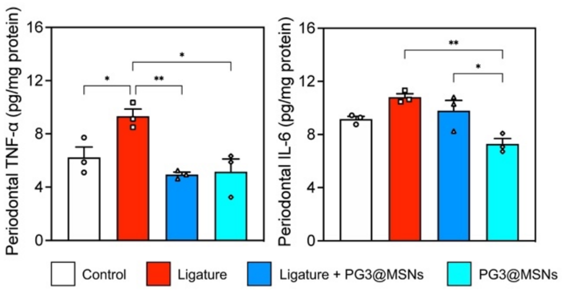


**Supplementary Fig. 10 Quantification of periodontal tissue TNF-α and IL-6 using ELISA.**

n = 3, Data are means ± SEM; differences were assessed by one-way analysis of variance and Tukey's multiple comparisons test. **P* < 0.05, ***P* < 0.01.

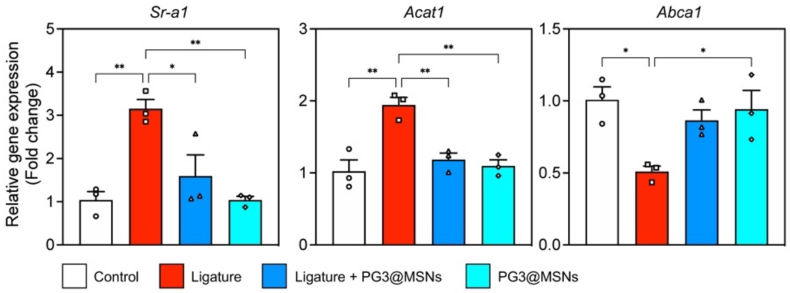


**Supplementary Fig. 11 Relative gene expression of *Sr-a1*, *Acat1*, and *Abca1* in thoracoabdominal aorta.**

n = 3, Data are means ± SEM; differences were assessed by one-way analysis of variance and Tukey's multiple comparisons test. **P* < 0.05, ***P* < 0.01.

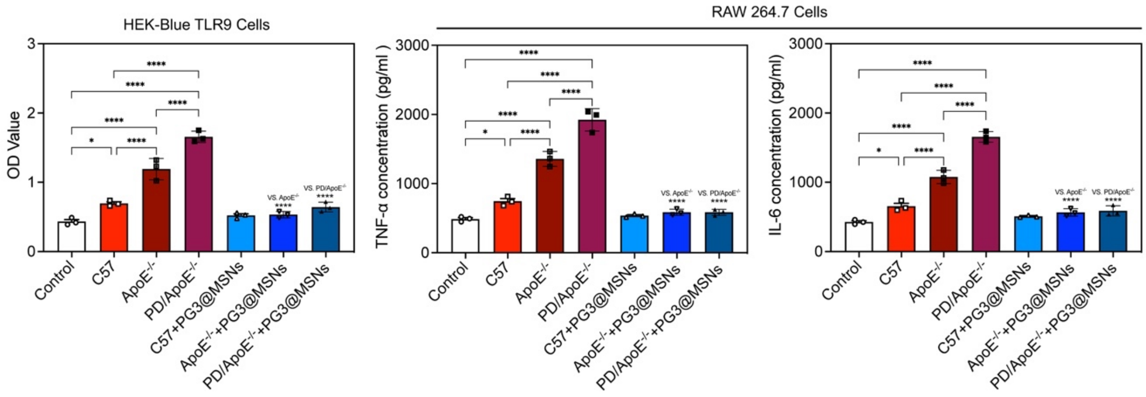


**Supplementary Fig. 12 In vivo anti-inflammatory stimulation experiment using body fluids.**

**a** Activation of HEK-Blue TLR9 reporter cells by plasma from 17-week-old C57BL/6 mice (C57), plasma from ApoE$^{-/-}$ mice in the Control group (ApoE$^{-/-}$), and plasma from ApoE$^{-/-}$ mice in the Ligature group (PD/ApoE$^{-/-}$) in the absence or presence of PG3@MSNs for 24 hours. The corresponding SEAP activity in supernatants from each group was determined with a QUANTI-Blue assay at $OD_{620}$.

**b** RAW 264.7 macrophages were stimulated with plasma C57, plasma ApoE$^{-/-}$, and plasma PD/ApoE$^{-/-}$ in the absence or presence of PG3@MSNs for 24 hours. Supernatants were assayed for TNF-α and IL-

6 by ELISA.

n = 3, Data are means ± SEM; differences were assessed by one-way analysis of variance and Tukey's multiple comparisons test. *$P < 0.05$, ****$P < 0.0001$.

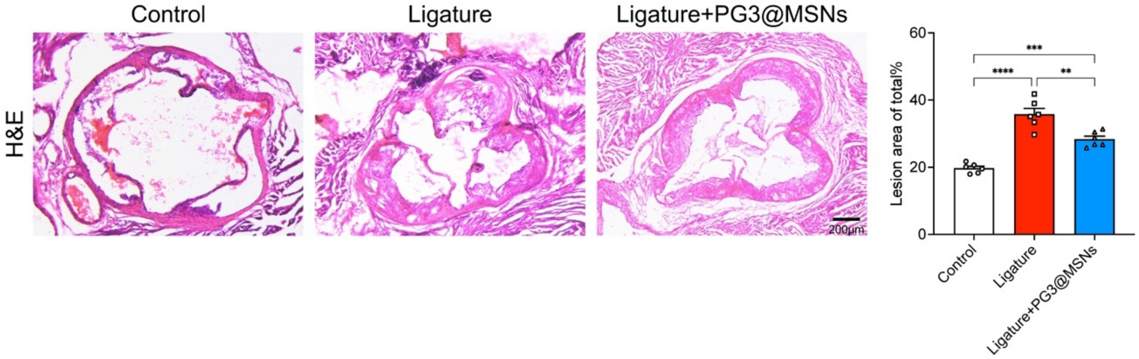


**Supplementary Fig. 13 H&E staining of aortic valve.**

Representative images of H&E staining of aortic valve (Scale bar, 200 μm) and quantification of plaque occupying the lumen area (n = 5).

Data are means ± SEM; statistical analysis was performed using one-way ANOVA with Tukey's post hoc test; **$P < 0.01$, ***$P < 0.001$, ****$P < 0.0001$.

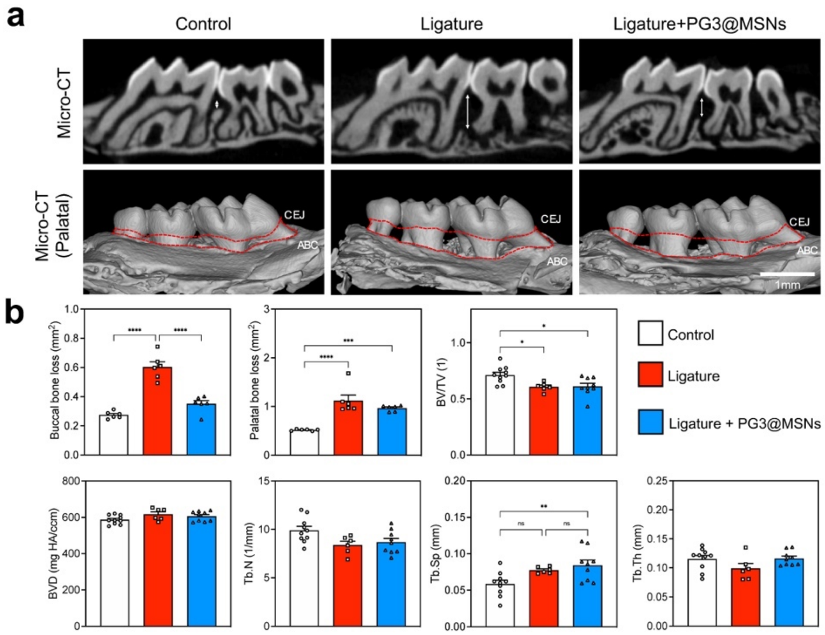


**Supplementary Fig. 14 Micro-CT of periodontal tissue.**

**a** Representative images of resliced view (top) and palatal view (bottom) in 3D reconstruction of mice maxilla. Scale bars, 1 mm.

**b** Quantification of buccal and palatal alveolar bone resorption (n = 8).

**c** Quantification of trabecular bone parameters in the furcation area of the maxillary second molar (n = 8). BV/TV, bone volume fraction. BVD, bone volume density. Tb.N, trabecular number. Tb.Sp, trabecular separation. Tb.Th, trabecular thickness.

Data are means ± SEM; statistical analysis was performed using one-way ANOVA with Tukey's post hoc test; *$P$ < 0.05, **$P$ < 0.01, ***$P$ < 0.001, ****$P$ < 0.0001.

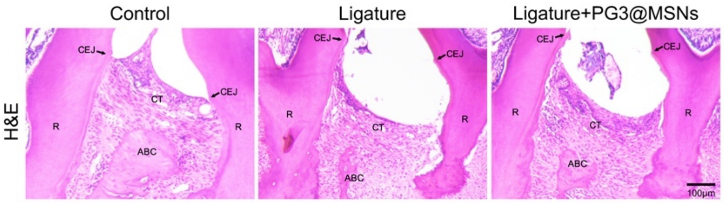


**Supplementary Fig. 15 H&E staining of periodontal tissue.**

Representative images of H&E staining of periodontal tissue. Scale bar, 200 μm.

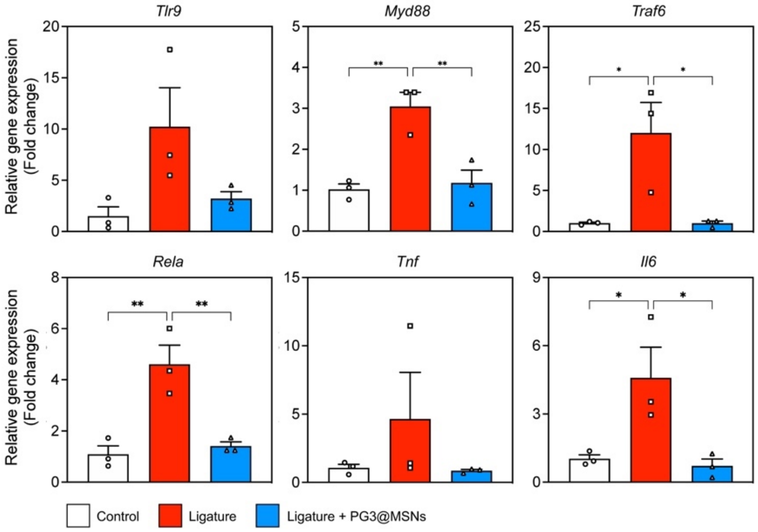


**Supplementary Fig. 16 Relative gene expression of *Tlr9*, *Myd88*, *Traf6*, *Rela*, *Tnf*, and *Il6* in thoracoabdominal aorta.**

n = 3, Data are means ± SEM; statistical analysis was performed using one-way ANOVA with Tukey's post hoc test; *$P$ < 0.05, **$P$ < 0.01.

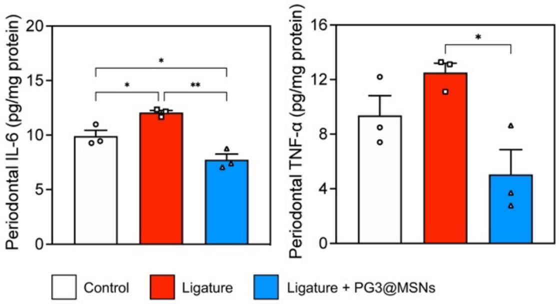


**Supplementary Fig. 17 Quantification of periodontal tissue TNF-α and IL-6 using ELISA.**

n = 3, Data are means ± SEM; statistical analysis was performed using one-way ANOVA with Tukey's post hoc test; *$P$ < 0.05, **$P$ < 0.01.

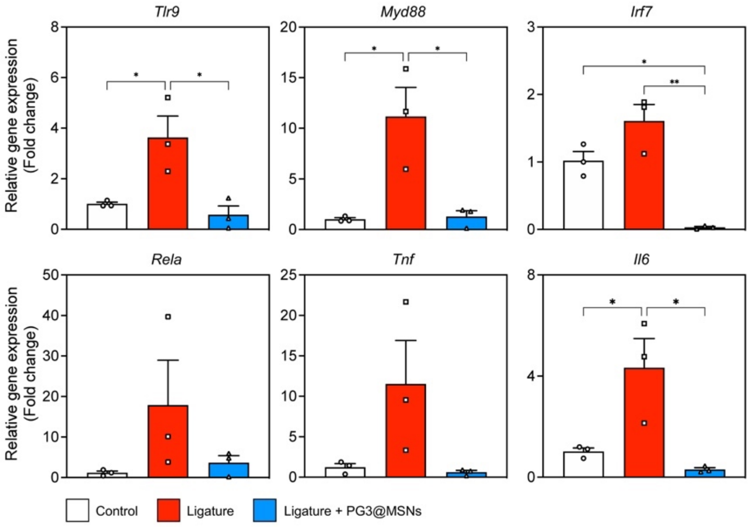


**Supplementary Fig. 18 Relative gene expression of *Tlr9*, *Myd88*, *Traf6*, *Rela*, *Tnf*, and *Il6* in periodontal tissue.**

n = 3, Data are means ± SEM; statistical analysis was performed using one-way ANOVA with Tukey's post hoc test; *$P < 0.05$, **$P < 0.01$.

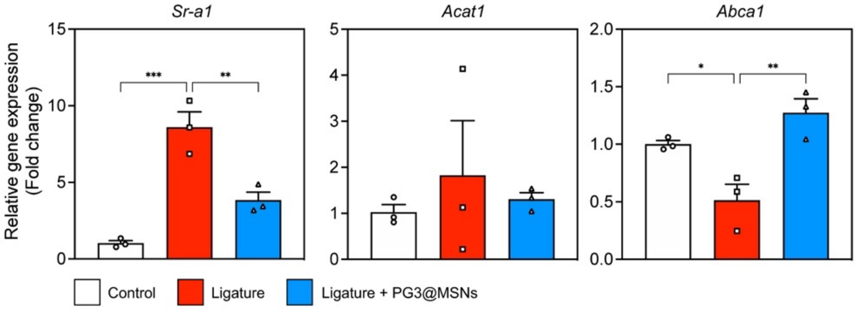


**Supplementary Fig. 19 Relative gene expression of *Sr-a1*, *Acat1*, and *Abca1* in thoracoabdominal aorta.**

n = 3, Data are means ± SEM; statistical analysis was performed using one-way ANOVA with Tukey's post hoc test; *$P < 0.05$, **$P < 0.01$, ***$P < 0.001$.

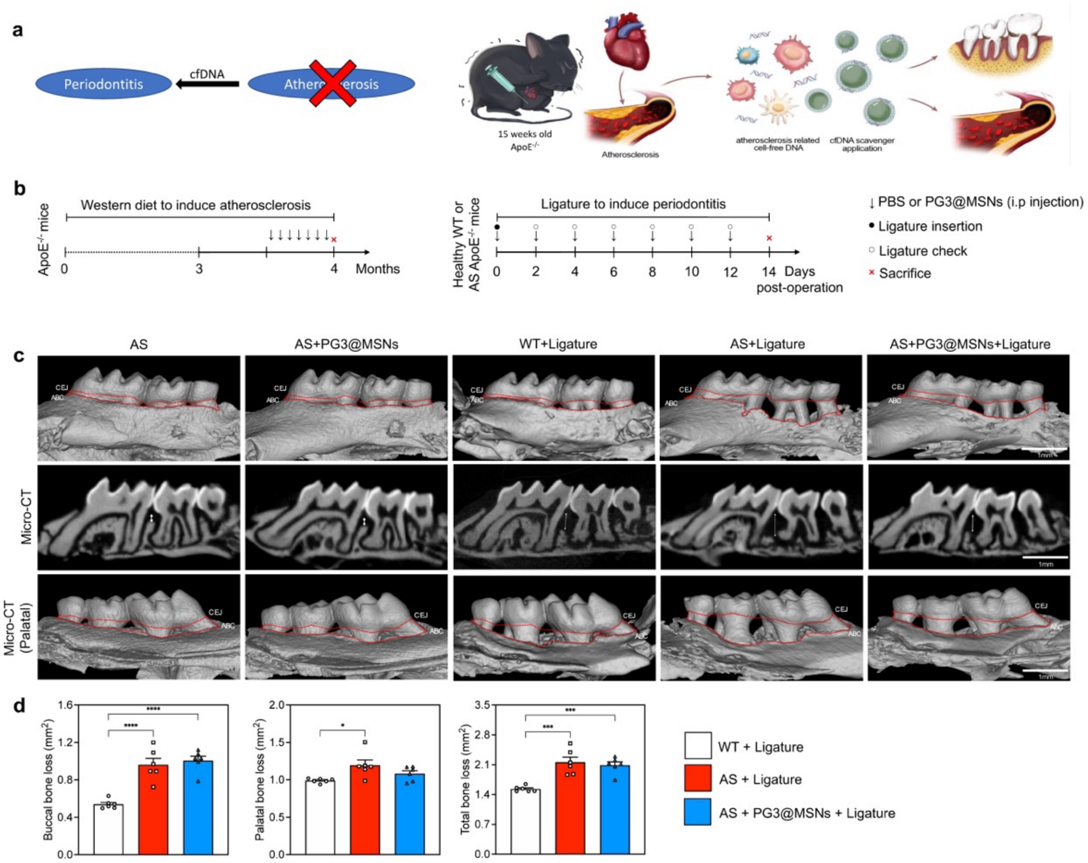


**Supplementary Fig. 20 Atherosclerosis deteriorated the experimental periodontitis and systematical administration of PG3@MSNs functioned little on alveolar bone loss.**

**a** Schematic illustration of the comorbidity relationship and the mechanism of PG3@MSNs systemically application.

**b** Experimental schedule of atherosclerosis induction (left) and periodontitis induction (right) with correspondent treatment in vivo study.

**c** Representative images of 3D reconstruction of mice maxilla. Scar bars, 1mm.

**d** Quantification of buccal, palatal, and total alveolar bone resorption in **c** (n = 3). Data are means ± SEM; statistical analysis was performed using one-way ANOVA with Tukey's post hoc test; $*P < 0.05$, $***P < 0.001$, $****P < 0.0001$.

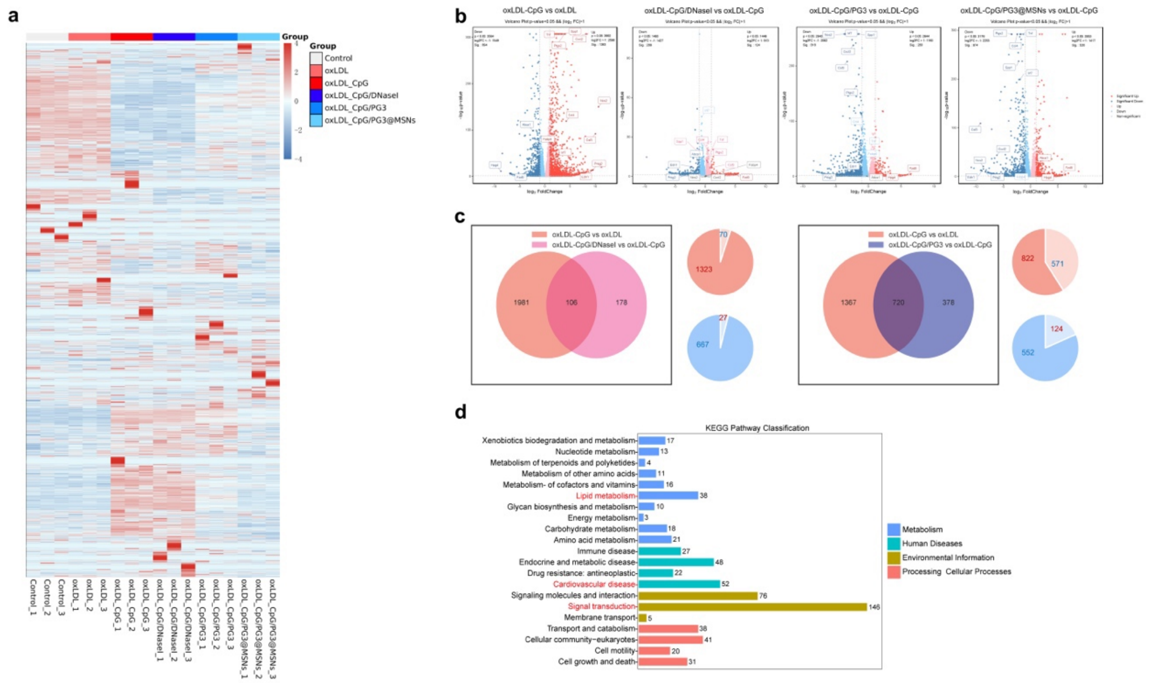


**Supplementary Fig. 21 Bulk RNA-seq profiling of cfDNA stimulated RAW 264.7 macrophages.**

**a** Heatmap showing genes expression profiling among RAW 264.7 macrophages with varied treatments (n = 3).

**b** Volcano plots illustrating the differential gene expression profiles of oxLDL-induced macrophages with or without cfDNA stimulation, as well as the changes in differentially expressed genes （DEGs） of macrophages treated with different materials.

**c** Venn diagram of the intersection of differentially expressed genes and pie charts indicating the number of DEGs regulated by respective materials.

**d** Classification of the enriched KEGG pathways of the intersection genes in **Fig. 5f**.

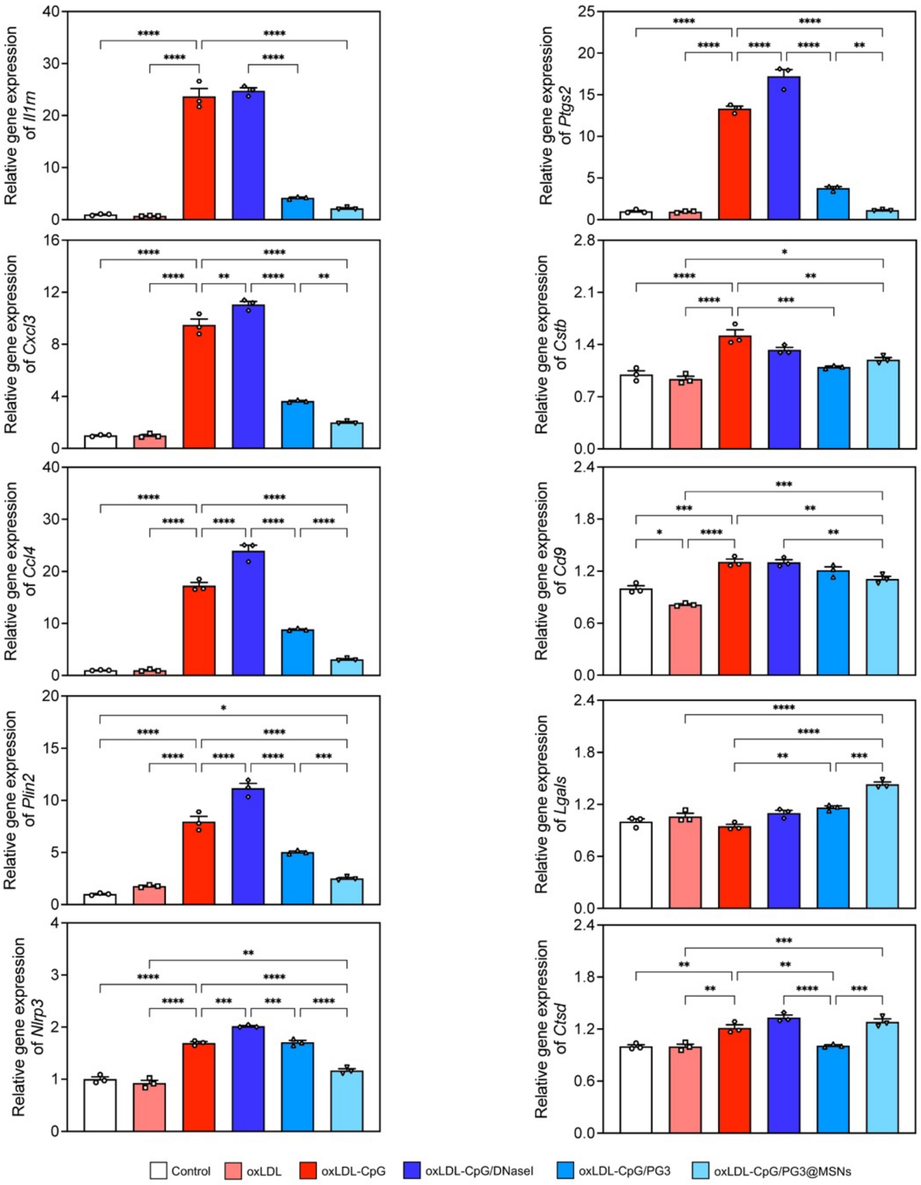


**Supplementary Fig. 22 Relative gene expression of inflammation- and foam cell-associated genes.** Relative gene expression of inflammation- and foam cell-associated genes in RAW 264.7 macrophages (n = 3). Data are means ± SEM; statistical analysis was performed using one-way ANOVA with Tukey's post hoc test; *$P < 0.05$, **$P < 0.01$, ***$P < 0.001$, ****$P < 0.0001$.

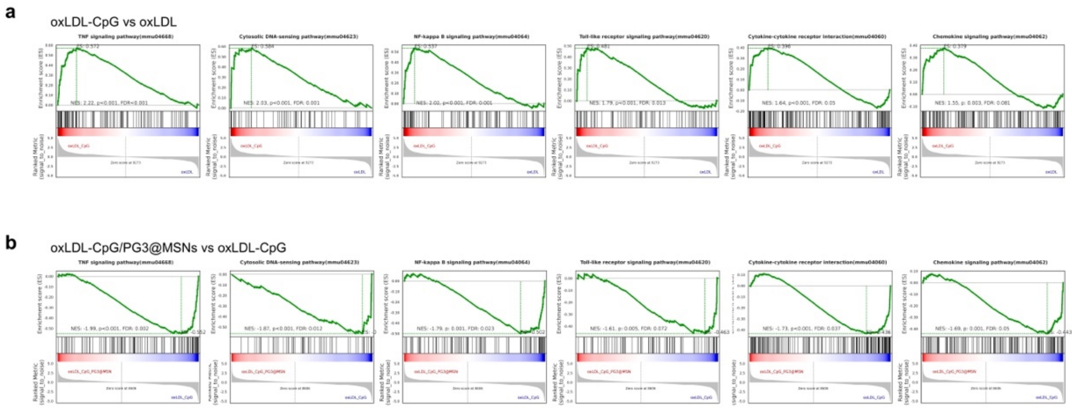


**Supplementary Fig. 23 Inflammation-related Gene set enrichment analysis.**

**a** cfDNA-induced changes of upregulation of the TNF, cytosolic DNA sensing, NF-κB, TLR, cytokine-cytokine receptor interaction and chemokine signaling pathways as assessed by Gene set enrichment analysis (GSEA) enrichment plots.

**b** Downregulation of the upregulated pathways mentioned in **a** by PG3@MSNs.

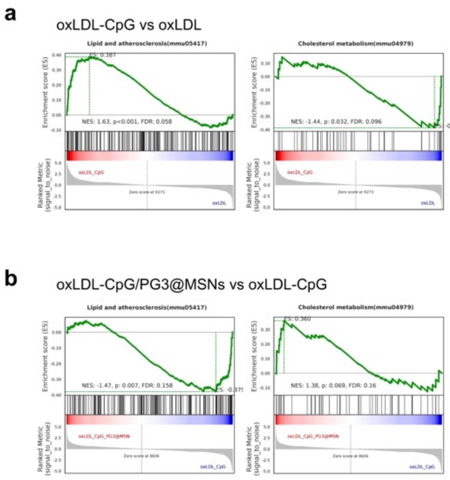


**Supplementary Fig. 24 Lipid metabolism-related Gene set enrichment analysis.**

**a** cfDNA-induced upregulation of the lipid and atherosclerosis pathway and downregulation of the cholesterol metabolism pathway as assessed by Gene set enrichment analysis (GSEA) enrichment plots.

**b** Reversion of the changed pathways mentioned in **a** by PG3@MSNs.

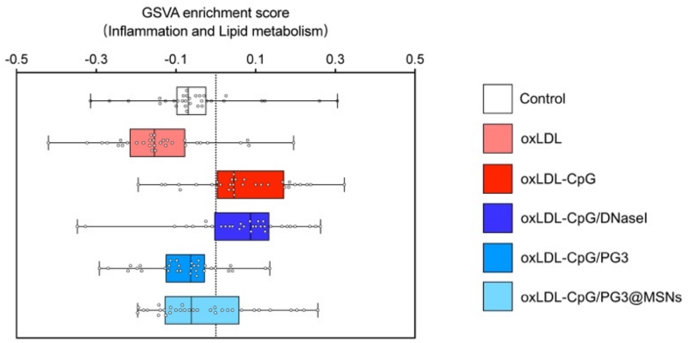


**Supplementary Fig. 25 Gene set variation analysis.**

GSVA score of inflammation- and lipid metabolism-related pathways of RAW 264.7 macrophages (n = 3). Data are means ± SEM.

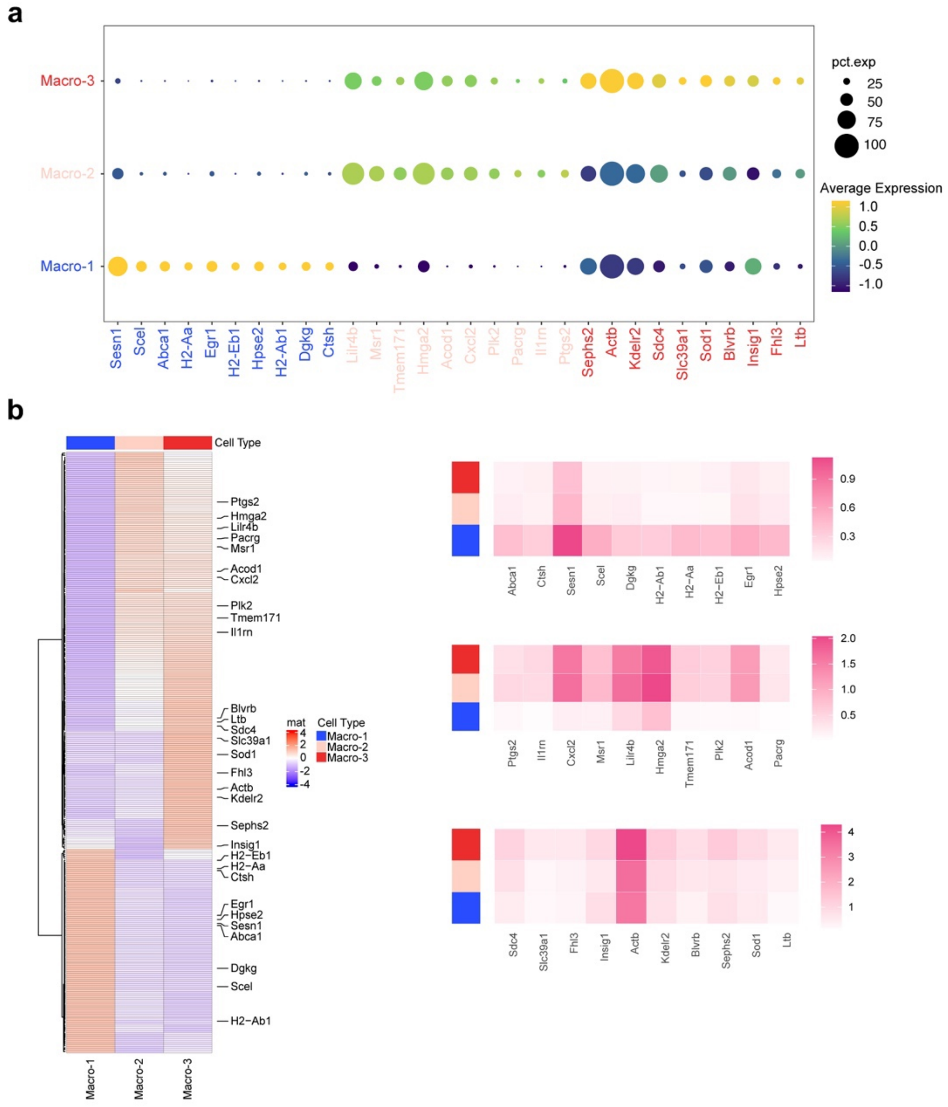


**Supplementary Fig. 26 Annotation of macrophage subcluster with marker genes.**

**a-b** Dot plot (**a**) and heatmaps (**b**) of marker genes in each macrophage subcluster.

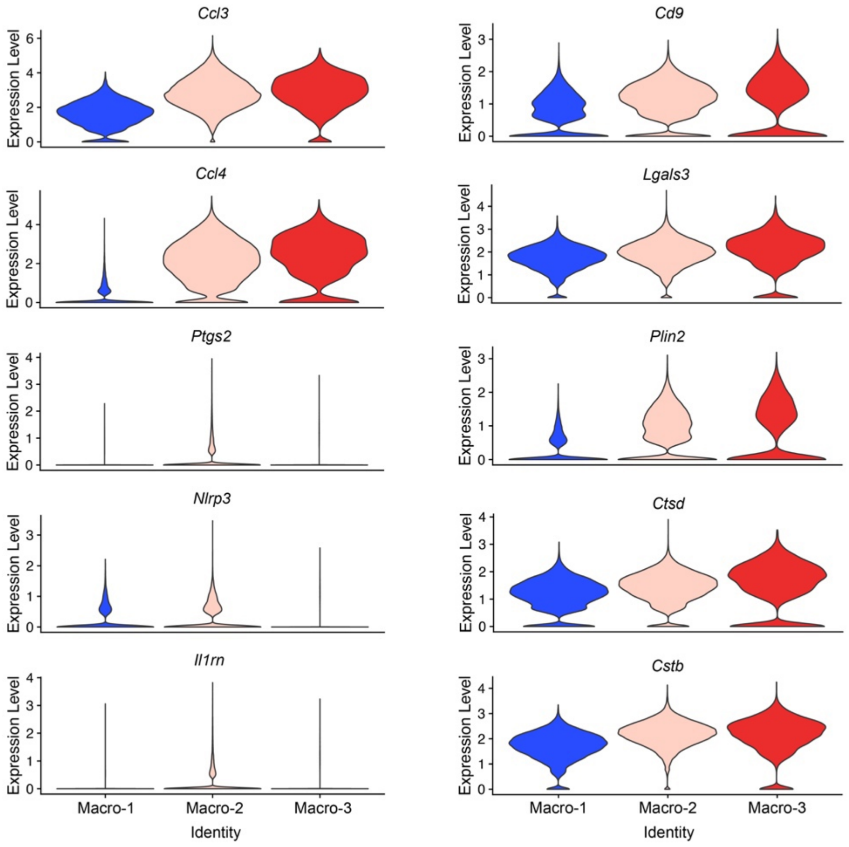


**Supplementary Fig. 27 Expression level of inflammation- and foam cell-related genes.**
Expression level of inflammation-related genes (left) and foam cell-related genes (right) in different macrophage subclusters.

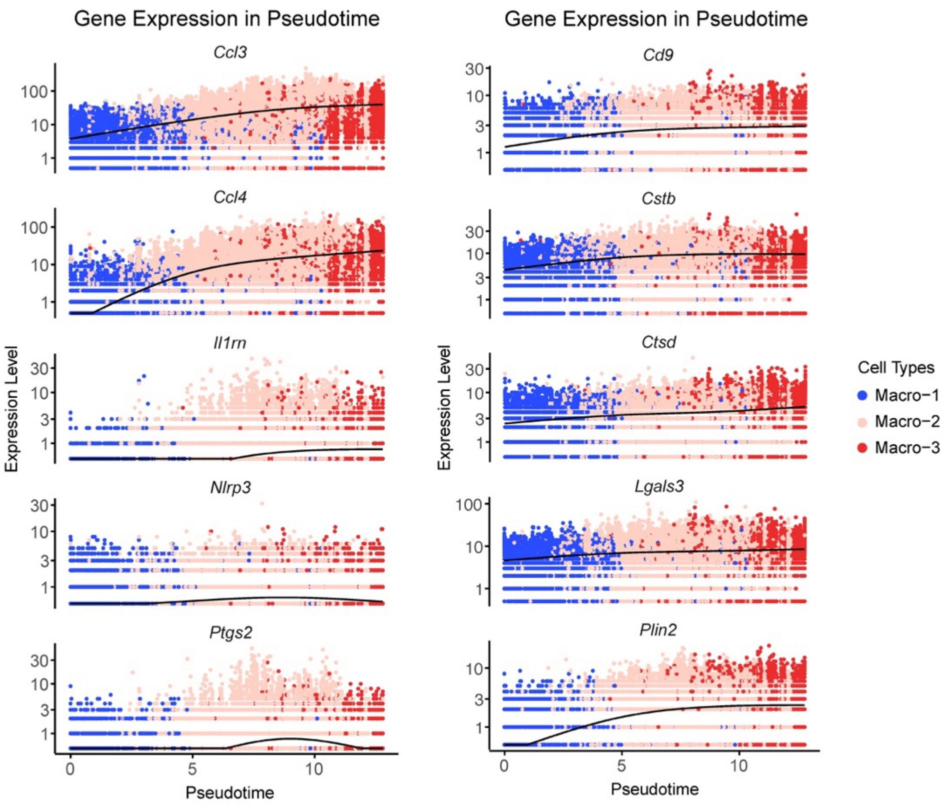


**Supplementary Fig. 28 Expression level of inflammation- and foam cell-related genes in pseudotime.**

Expression level of inflammation-related genes (left) and foam cell-related genes (right) in different macrophage subclusters along the differentiation trajectory in pseudotime.
